# Predictions and experimental tests of a new biophysical model of the mammalian respiratory oscillator

**DOI:** 10.1101/2021.10.29.466442

**Authors:** Ryan S. Phillips, Hidehiko Koizumi, Yaroslav I. Molkov, Jonathan E. Rubin, Jeffrey C. Smith

## Abstract

Previously our computational modeling studies (Phillips et al., 2019) proposed that neuronal persistent sodium current (I_NaP_) and calcium-activated non-selective cation current (I_CAN_) are key biophysical factors that, respectively, generate inspiratory rhythm and burst pattern in the mammalian preBötzinger complex (preBötC) respiratory oscillator. Here, we experimentally tested and confirmed three predictions of the model from new simulations concerning the roles of I_NaP_ and I_CAN_: (1) I_NaP_ and I_CAN_ blockade have opposite effects on the relationship between network excitability and preBötC rhythmic activity; (2) I_NaP_ is essential for preBötC rhythmogenesis; (3) I_CAN_ is essential for generating the amplitude of rhythmic output but not rhythm generation. These predictions were confirmed via optogenetic manipulations of preBötC network excitability during graded I_NaP_ or I_CAN_ blockade by pharmacological manipulations in neonatal mouse slices *in vitro*. Our results support and advance the hypothesis that I_NaP_ and I_CAN_ mechanistically underlie rhythm and inspiratory burst pattern generation, respectively, in the isolated preBötC.

## Introduction

The cellular and circuit biophysical mechanisms generating the breathing rhythm critical for life in mammals have been under investigation for decades without clear resolution (Richter and Smith, 2014; Del Negro, Funk and Feldman, 2018; Ramirez and Baertsch, 2018). Biophysical mechanisms operating in the mammalian respiratory oscillator within the brainstem preBötzinger complex (preBötC), which contains essential rhythmogenic neuronal circuits, have recently been re-examined with a new computational model that builds on previous models and proposes distinct functional roles for cellular-level sodium- and calcium-based biophysical mechanisms that have been of intense experimental and theoretical interest for understanding the operation of preBötC circuits (Phillips *et al*., 2019). These mechanisms rely on a slowly inactivating persistent sodium current (I_NaP_) and a calcium-activated non-selective cation current (I_CAN_) mediated by transient receptor potential M4 (TRPM4) channels coupled to intracellular calcium dynamics. The model proposes how these cellular-level mechanisms operating in excitatory circuits can underlie two critical functions of preBötC circuits: generation of the inspiratory rhythm, and regulation of the amplitude of inspiratory population activity. In essence, the model advances the concepts that (1) a subset of excitatory circuit neurons whose rhythmic bursting is critically dependent on I_NaP_ forms an excitatory neuronal kernel for rhythm generation, and (2) excitatory synaptic drive from the rhythmogenic kernel population is critically amplified by I_CAN_ activation in recruited and interconnected preBötC excitatory neurons to generate the amplitude of population activity necessary to drive downstream excitatory circuits to produce inspiratory motor output.

While these concepts have some experimental support (see discussions and model-experimental data comparisons in (Phillips *et al*., 2019)), the mechanisms incorporated in the model and their predicted effects on circuit behavior require more extensive experimental testing. Indeed, the basic rhythmogenic I_NaP_-based biophysical mechanism represented by the model remains controversial (Del Negro, Funk and Feldman, 2018). The concept that activation of I_CAN_ is largely due to synaptically-activated sources of neuronal Ca^2+^ flux, such that I_CAN_ contributes to the excitatory inspiratory drive potential and regulates inspiratory burst amplitude by mirroring the excitatory synaptic current, also requires additional experimental support. In the present modeling and experimental study, to advance our previous studies (Phillips *et al*., 2019), we have performed experimental tests on the rhythmically active *in vitro* network in slices from transgenic mice using a combination of electrophysiological analyses, pharmacological perturbations of I_NaP_ or I_CAN_, and optogenetic manipulations of the preBötC excitatory (glutamatergic) population involved. Our new model simulation-experimental data comparisons provide further evidence for the predictive power and validity of the model, while also indicating some limitations.

## Results

### Model concepts and predictions

The following features of the model outputs from new simulations (Figure 1) formed the basis for the design of experiments for model simulation-experimental data comparisons in the current study. Simulations were performed with the model specified in Phillips *et al*. (2019), with modifications as described in Materials and Methods.

**Figure 1.**
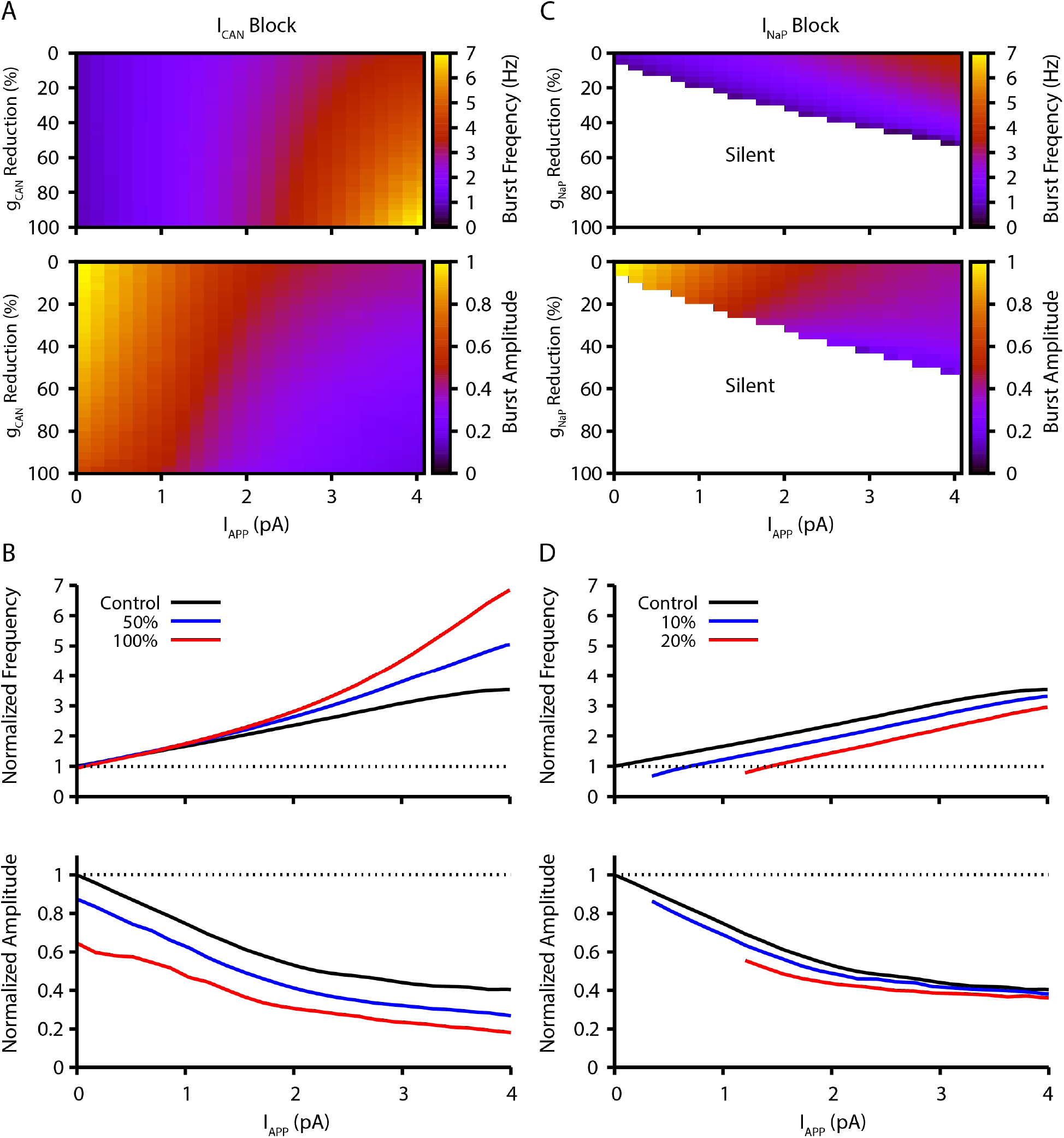
Model simulation predictions for the relationships between applied current (I_App_) and population burst frequency and amplitude of synaptically-coupled excitatory neurons (N = 100) incorporating I_NaP_ and I_CAN_ over a range of conductances (g_CAN_, g_NaP_). Pharmacological block of I_NaP_ and I_CAN_ is simulated by a percent reduction of their respective conductances. (**A & B**) Model parameter space plots color-coded from simulations show effects on frequency (upper panel in **A**) and amplitude (lower panel in **A**) of reductions in g_CAN_, and in **B** at fixed levels of reductions (50%, 100%) in g_CAN_ from initial control values over a wide range of I_App_. (**C & D**) Parameter space plots showing effects in **C** of reductions in g_NaP_ on frequency and amplitude of model neuronal population activity, and in **D** at fixed levels of reduction (10%, 20%) in g_NaP_ from control. Color scale bars for values of burst frequency and amplitude are at right of plots in **A** and **C**. Frequencies and amplitudes of population activity in **C** and **D** are normalized to control values.

(1) The inspiratory rhythm *in vitro* is generated by a functionally distinct subpopulation of preBötC excitatory neurons with subthreshold-activating I_NaP_ that confers voltage-dependent rhythmic burst frequency control, providing a mechanism for frequency tuning by the regulation of baseline membrane potentials. Progressively depolarizing/hyperpolarizing neurons across the population by varying an applied current (I_App_) increases/decreases population burst frequency over a wide dynamic range defined by the frequency tuning curve for the population (i.e., the relationship between applied current/baseline membrane potential and network bursting frequency).

(2) Reducing neuronal persistent sodium conductance (g_NaP_) decreases the population-level bursting frequency and alters the voltage-dependent rhythmogenic behavior with a shift in the frequency tuning curve such that the frequency dynamic range is reduced, consistent with previous simulations (Phillips and Rubin, 2019). Population amplitude is only slightly reduced by decreases in g_NaP_. Rhythm generation can be terminated at a sufficiently low level of g_NaP_ under baseline conditions of network excitability *in vitro*.

(3) Reducing g_CAN_ strongly decreases population activity amplitude and, in contrast to reducing g_NaP_, has little effect on population bursting frequency under baseline conditions, but strongly augments bursting frequency due to a shift in the frequency tuning curve that extends the frequency range at higher levels of population depolarization.

Model simulations showing the relations between I_App_, population burst frequency, and burst amplitude at different levels of g_NaP_ and g_CAN_ are displayed in Figure 1, which illustrate these basic predictions of the model. These contrasting predictions for effects of g_NaP_ or g_CAN_ on network dynamics provide a clear basis for experimental testing of the model.

### Experimental design and results

For comparisons with modeling results, we used electrophysiological recording from the preBötC inspiratory population and the hypoglossal (XII) motor output, in conjunction with pharmacological and optogenetic manipulations of preBötC glutamatergic neurons in *in vitro* slices. Slices were prepared from transgenic mice that selectively express Channelrhodopsin-2 (ChR2) in glutamatergic neurons by Cre-dependent targeting via the vesicular glutamate transporter-type 2 (VgluT2) promoter (Figure 2). ChR2-based photostimulation of preBötC glutamatergic neurons bilaterally enabled graded control of the baseline depolarization of this excitatory population to define the frequency tuning curves and associated population bursting amplitudes at different levels of pharmacological block of g_NaP_ or g_CAN_.

**Figure 2.**
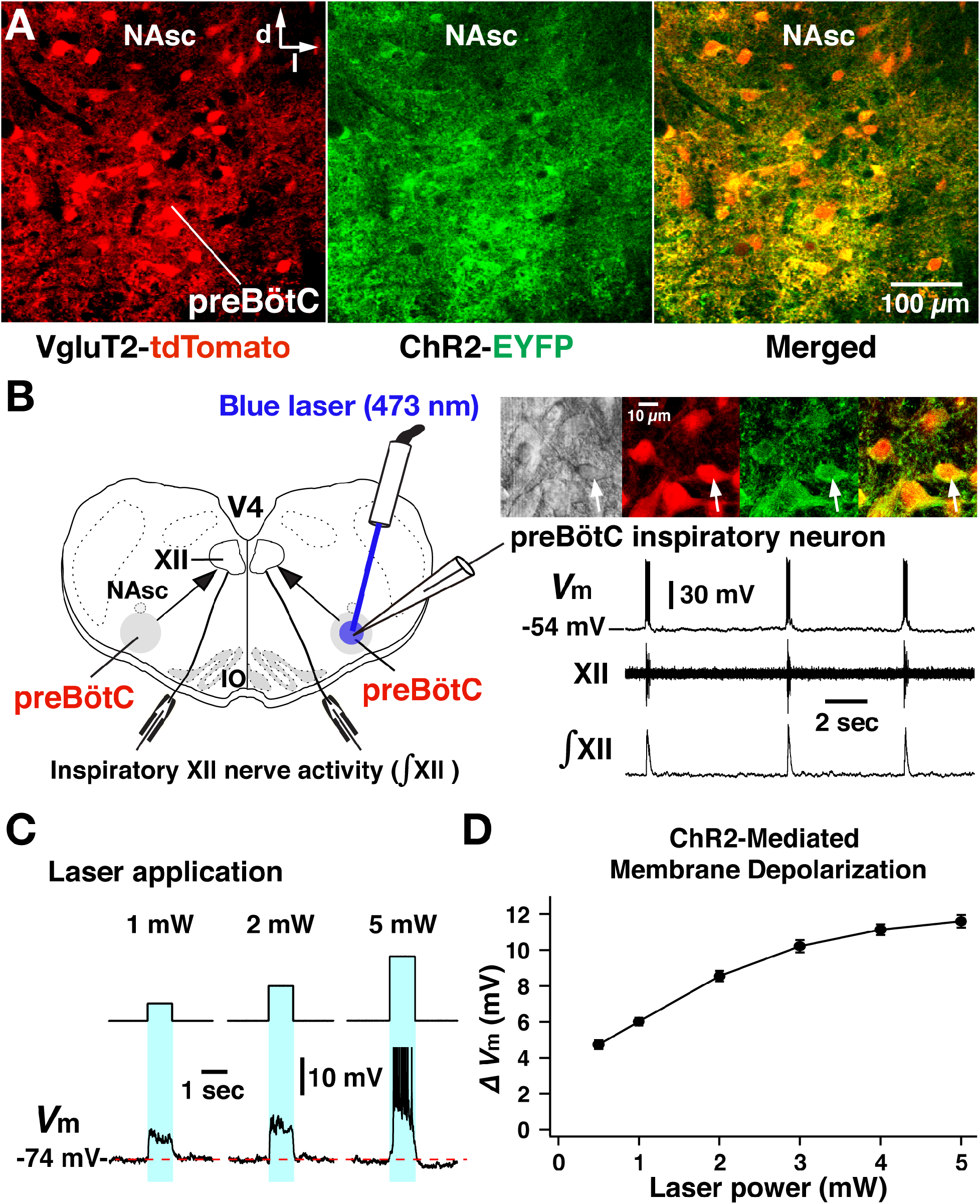
ChR2-mediated membrane depolarization of preBötC VgluT2-positive inspiratory neurons *in vitro*. (**A**) Two-photon microscopy single optical plane live images of the preBötC subregion in an *in vitro* neonatal medullary slice preparation from the VgluT2-tdTomato-ChR2-EYFP transgenic mouse line, illustrating tdTomato-labeled VgluT2-positive neurons distributed in the preBötC and adjacent regions (red) and expression of ChR2-EYFP (green) in somal membranes and neuronal processes of VgluT2-tdTomato neurons, as seen in the merged image. Abbreviations: d, dorsal; l, lateral; NAsc, semi-compact subdivision of nucleus ambiguous (NA). (**B**) Overview of experimental *in vitro* rhythmic slice preparation from a neonatal VgluT2-tdTomato-ChR2 transgenic mouse showing whole-cell patch-clamp recording from a functionally identified preBötC inspiratory VgluT2-positive neuron with unilateral preBötC laser illumination (0.5 – 5 mW) to test for neuronal expression of ChR2, and suction-electrode extracellular recordings from hypoglossal (XII) nerves to monitor inspiratory activity. Regional photostimulation was accomplished with a 100 μm diameter optical cannula positioned at the slice surface above the preBötC region. Abbreviations: V4, fourth ventricle; IO, inferior olivary nucleus. Upper right in **B**: two-photon single optical plane images showing an imaged preBötC inspiratory neuron (arrow) targeted for whole-cell recording. From left, Dodt structural image, VgluT2-Cre driven tdTomato labeling, ChR2-EYFP expression and merged image that confirms co-expression of tdTomato and ChR2-EYFP. The lower right traces are current-clamp recordings from the VgluT2-positive preBötC neuron shown in the above images, illustrating inspiratory spikes/bursts synchronized with integrated inspiratory XII nerve activity (∫XII). (**C**) The membrane potential (*V*_m_) of the neuron shown in **B** was depolarized during photostimulation by ~6.0 mV at 1 mW, ~8.5 mV at 2 mW and ~11.5 mV at 5 mW of laser power (spikes are truncated). The neuron was hyperpolarized from resting baseline potential to −74 mV by applying constant current in this example to reveal the magnitude of the light-induced membrane depolarization. Photostimulation was performed in the interval between inspiratory population bursts. (**D**) Summary data (n = 12 neurons from 4 slice preparations, mean ± SEM) showing the laser-power dependent, ChR2-mediated neuronal membrane depolarization from baseline (ΔV_M_) of VgluT2-positive preBötC inspiratory neurons.

#### Cre-dependent targeting of preBötC glutamatergic neurons for photostimulation experiments

For optogenetic photostimulation, we established and validated a triple transgenic VgluT2-tdTomato-ChR2-EYFP mouse line, produced by crossing a previously validated VgluT2-Cre-tdTomato mouse (Koizumi *et al*., 2016) with a Cre-dependent ChR2-EYFP mouse strain. We verified Cre-driven ChR2-EYFP expression in VgluT2-positive tdTomato labeled neurons within the preBötC region by two-photon microscopy in live neonatal mouse medullary slice (n = 6) preparations *in vitro*, documenting heavy expression of the ChR2-EYFP fusion protein in processes and somal membranes of VgluT2-tdTomato labeled neurons (Figure 2A). We also quantified by whole-cell patch-clamp recordings the photostimulation-induced depolarization of rhythmically active, ChR2-expressing inspiratory glutamatergic neurons (n = 12 neurons from 4 slice preparations from VgluT2-tdTomato-ChR2-EYFP mouse line) at different levels of photostimulation (473 nm of variable laser power ranging from 0.5 to 5 mW) in rhythmically active *in vitro* neonatal medullary slice preparations. These *in vitro* slices effectively isolate the bilateral preBötC along with hypoglossal (XII) motoneurons and nerves to monitor inspiratory XII activity (Koizumi *et al*., 2016; Koizumi *et al*., 2013), allowing laser illumination of the preBötC region unilaterally or bilaterally as well as whole-cell patch-clamp recordings from rhythmically-active inspiratory preBötC neurons during laser illumination (Figure 2B). We performed targeted current-clamp recordings from rhythmic tdTomato-labeled VgluT2-positive neurons, in which co-expression of ChR2-EYFP fusion protein in the cell membranes was confirmed with two-photon, single optical plane live images (Figure 2B, right). Representative examples of ChR2-mediated membrane depolarization (Figure 2C) shows that the membrane potential of a functionally identified VgluT2-positive preBötC inspiratory neuron was depolarized by ~6.0 mV at 1 mW, ~8.5 mV at 2 mW and ~11.5 mV at 5 mW laser illumination. The light-induced depolarization had fast kinetics, occurring within ~50 ms, and recovery within ~150 ms after terminating illumination. Summary data (n = 12 neurons, mean ± SEM, Figure 2D) illustrates the laser-power dependent ChR2-mediated depolarization of VgluT2-positive inspiratory preBötC neurons (4.73 ± 0.24 mV at 0.5 mW, 6.01 ± 0.21 mV at 1 mW, 8.55 ± 0.29 mV at 2 mW, 10.21 ± 0.34 mV at 3 mW, 11.13 ± 0.3 mV at 4 mW, and 11.6 ± 0.35 mV at 5 mW) (membrane depolarizations are significantly different, p<0.01, in all cases, one sample *t*-test), enabling us to define the relation between laser power and ChR2-mediated membrane depolarization for the inspiratory glutamatergic neuron population.

#### Perturbations of inspiratory rhythm and burst amplitude by photostimulation of glutamatergic neurons within the preBötC region

We performed site-specific bilateral photostimulations of the preBötC VgluT2-positive neuron population (Figure 3A) by sustained laser illumination (20 ms pulses at 20 Hz, 473 nm, 30 - 90 sec pulse train durations) at graded laser powers between 0.25 – 2.0 mW (n = 31 slice preparations). To systematically analyze relations between inspiratory burst frequency/burst amplitude vs laser power, we applied single epochs of laser illumination and allowed for recovery from each epoch before changing the laser power [representative examples with simultaneous recordings from preBötC and XII are shown in Figure 3B, and summary data (mean ± SEM) is shown in Figure 3C]. Bilateral photostimulation (0.25 to 2 mW) of the preBötC caused rapid and reversible increases of inspiratory burst frequency in a laser-power dependent manner (85 ± 7% increase at 0.25 mW, 174 ± 7% at 0.5 mW, 262 ± 6% at 1 mW, 326 ± 6% at 2 mW, one sample *t*-test, p<0.01 in all cases). After termination of laser illumination, an inhibition of rhythmicity followed, and subsequently recovered to the control frequency within ~1 to 3 min. Concurrently, there was a significant laser power-dependent decrease of burst amplitude of inspiratory preBötC population and inspiratory XII activity during photostimulation (8 ± 2% decrease at 0.25 mW, 15 ± 2% at 0.5 mW, 25 ± 3% at 1 mW, 44 ± 4% at 2 mW, p<0.01 in all cases by Wilcoxon signed rank test). Bilateral photostimulation also induced tonic activity in preBötC population recordings, which was of higher amplitude at higher laser power, but did not consistently induce tonic activities in XII nerve output. Otherwise, XII burst amplitude and frequency always directionally mirrored preBötC glutamatergic population activity.

**Figure 3.**
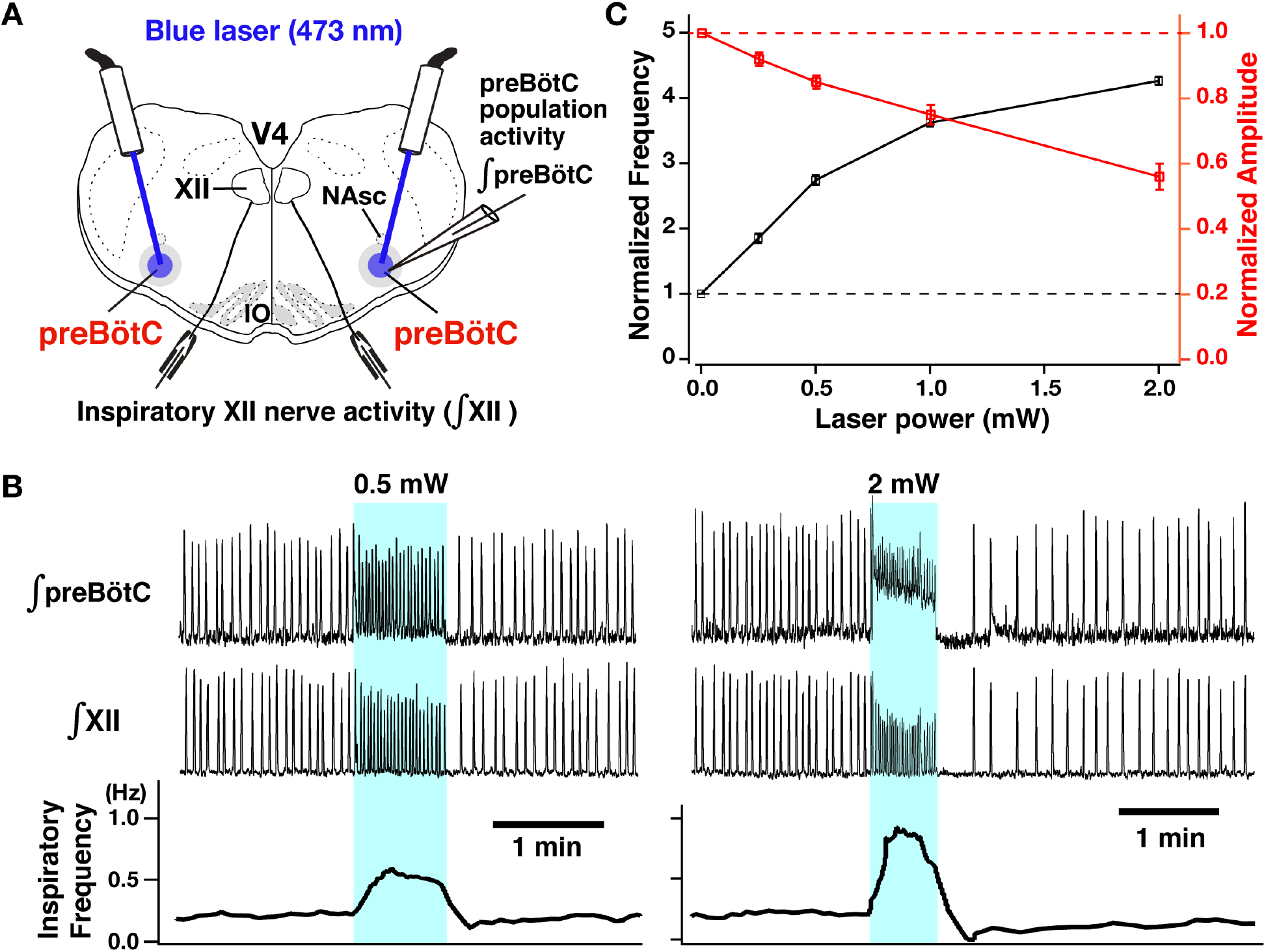
Photostimulation of the bilateral preBötC VgluT2-positive neuron population caused laser power dependent increases of inspiratory burst frequency and decreases of burst amplitude. (**A**) Overview of experimental *in vitro* rhythmically active slice preparation from neonatal VgluT2-tdTomato-ChR2 transgenic mouse with macro-patch electrodes on the preBötC region for recording preBötC population activity and suction-electrodes on hypoglossal (XII) nerves to monitor inspiratory motor activity during bilateral preBötC laser illumination (0.25 – 2 mW) with optical cannula (100 μm diameter) positioned obliquely to illuminate the preBötC region. Abbreviations: NAsc, nucleus ambiguus semi-compact subdivision; V4, fourth ventricle; IO, inferior olivary nucleus. (**B**) Representative examples of epochs of bilateral preBötC laser illumination and effects on inspiratory burst frequency and burst amplitude. The upper traces show the integrated macro-patch recordings from the preBötC inspiratory neuron population (∫preBötC), middle traces show the integrated inspiratory XII activity (∫XII), and the bottom traces show the inspiratory burst frequency (time-based moving median in a 10 s window). The sustained laser illumination (473 nm) with the laser intensity is given by blue shading. Low-intensity illumination (0.5 mW) caused significant increase (~149% in this example) of inspiratory burst frequency and decrease of inspiratory burst amplitude of both preBötC and XII population activity (~22% and ~21%, respectively, left panel). Higher intensity laser illumination (2 mW) (right panel) caused larger increase (~295%) of inspiratory frequency and decrease of burst amplitude of inspiratory preBötC and XII activity (~65% and ~44%, respectively). Also note that photostimulation induced tonic activity (indicated by baseline shift) in ∫preBötC population activity recordings. (**C**) Summary data of relations between inspiratory burst frequency and amplitude vs photostimulation laser power (n = 31, mean ± SEM).

These results demonstrate that preBötC glutamatergic neurons play an important role in regulating the inspiratory rhythm with a membrane voltage-dependent frequency control mechanism in neonatal medullary slices *in vitro*. We note that in control experiments with rhythmic slice preparations from the VgluT2-tdTomato, non-ChR2-expressing transgenic mice (n = 6), we tested for photo-induced perturbations of inspiratory burst frequency and amplitude, and confirmed that there were no significant perturbations of frequency and amplitude as a function of laser power (0.25-2 mW; Wilcoxon signed rank test for frequency and burst amplitude, p = 0.249 and 0.883, respectively).

#### Regulation of burst frequency and amplitude by I_NaP_ in preBötC glutamatergic neurons analyzed with combined optogenetic and pharmacological manipulations

After defining the voltage-dependent regulation of burst frequency and amplitude as described above, we investigated how pharmacological block of I_NaP_ in preBötC glutamatergic neurons affects this voltage-dependent control in rhythmically active *in vitro* medullary slice preparations from the VgluT2-tdTomato-ChR2 mouse line. We first analyzed the pharmacological profile of block of I_NaP_ in preBötC inspiratory glutamatergic neurons with the I_NaP_ blockers tetrodotoxin citrate (TTX) at very low concentrations (5 – 20 nM, n=8) and also riluzole (5 – 20 μM, n=6) bath-applied in the neonatal mouse medullary slice preparations as we have previously done for neonatal rat preBötC inspiratory neurons, but now analyzed for genetically identified mouse glutamatergic neurons. Whole-cell voltage-clamp recording from optically identified VgluT2-tdTomato-expressing inspiratory neurons was used to obtain subthreshold current-voltage (I-V) relationships by applying slow voltage ramps (30 mV/s; −100 to +10 mV) and we computed TTX- and riluzole-sensitive I_NaP_ by subtracting I-V curves obtained before and after block of I_NaP_ with TTX or riluzole (Figure S1) (Koizumi and Smith, 2008). Both TTX and riluzole attenuated the peak I_NaP_ amplitude (measured at −40 mV) in a dose-dependent manner. TTX reduced peak I_NaP_ by 36-63% at 5 nM, by 74-94% at 10 nM, and completely blocks I_NaP_ at 20 nM. Riluzole reduced peak I_NaP_ by 36-66% at 5 μM, by 82-95% at 10 μM, and completely blocks I_NaP_ at 20 μM. These pharmacological profiles are comparable to the data obtained in neonatal rat preBötC inspiratory neurons in our previous *in vitro* studies (Koizumi and Smith, 2008). We note that these pharmacological profiles were obtained for neurons recorded at depths up to 50-75 μm in the slices; neurons located deeper in the slice tissue may exhibit less reduction of I_NaP_ at the time points when these more superficial neurons were analyzed due to tissue penetration-related, spatiotemporal non-uniformity of I_NaP_ attenuation.

We then comparatively analyzed control of XII inspiratory burst frequency and amplitude by bilateral preBötC photostimulation (473 nm, 20 Hz pulses, 30 – 90 sec. trains of sustained photostimulation as described above) under I_NaP_ pharmacological blockade. As shown in Figures 4A and 4B, partial block of I_NaP_ (5 – 10 nM TTX, or 5 – 10 μM riluzole) gradually reduced XII inspiratory burst frequency, slightly reduced amplitude, and terminated inspiratory rhythmic bursting in almost all cases (n=6/7 at 5 nM TTX, n=8/8 at 10nM TTX, n=6/7 at 5 μM riluzole, n=8/8 at 10 μM riluzole). In the cases where rhythmic bursting was terminated, after 30 min. we again bilaterally photostimulated the preBötC, which restored rhythmic bursting with voltage-dependent frequency control (0.25 - 2.0 mW) in all cases (n=14/14 with TTX, n=14/14 with riluzole). The summary data (n=6 slice preparations for TTX 5nM, n=8 for TTX 10 nM, n=6 for riluzole 5 μM, n=8 for riluzole 10 μM, mean ± SEM) in Figure 4C (left) shows the frequency tuning curves with significant increases in burst frequency as a function of laser power (one sample *t*-test, p<0.01 in all cases except p<0.05 at 0.25 mW with 10 nM TTX and at 0.25 mW with 10 μM riluzole). The tuning curves are significantly shifted downward relative to control conditions (TTX 5 nM vs control, TTX 10 nM vs control, riluzole 5 μM vs control, riluzole 10 μM vs control; in all cases, p<0.01 by Wilcoxon matched pair signed rank test), and more downwardly-shifted with higher drug concentrations (Wilcoxon signed rank test, TTX 5 nM vs TTX 10 nM, p<0.01; riluzole 5 μM vs riluzole 10 μM, p = 0.018). The frequency-laser power relations are not significantly different for TTX and riluzole (Wilcoxon signed rank test, p = 0.101 for TTX 5 nM vs riluzole 5 μM; p = 0.431 for TTX 10 nM vs riluzole 10 μM).

**Figure 4.**
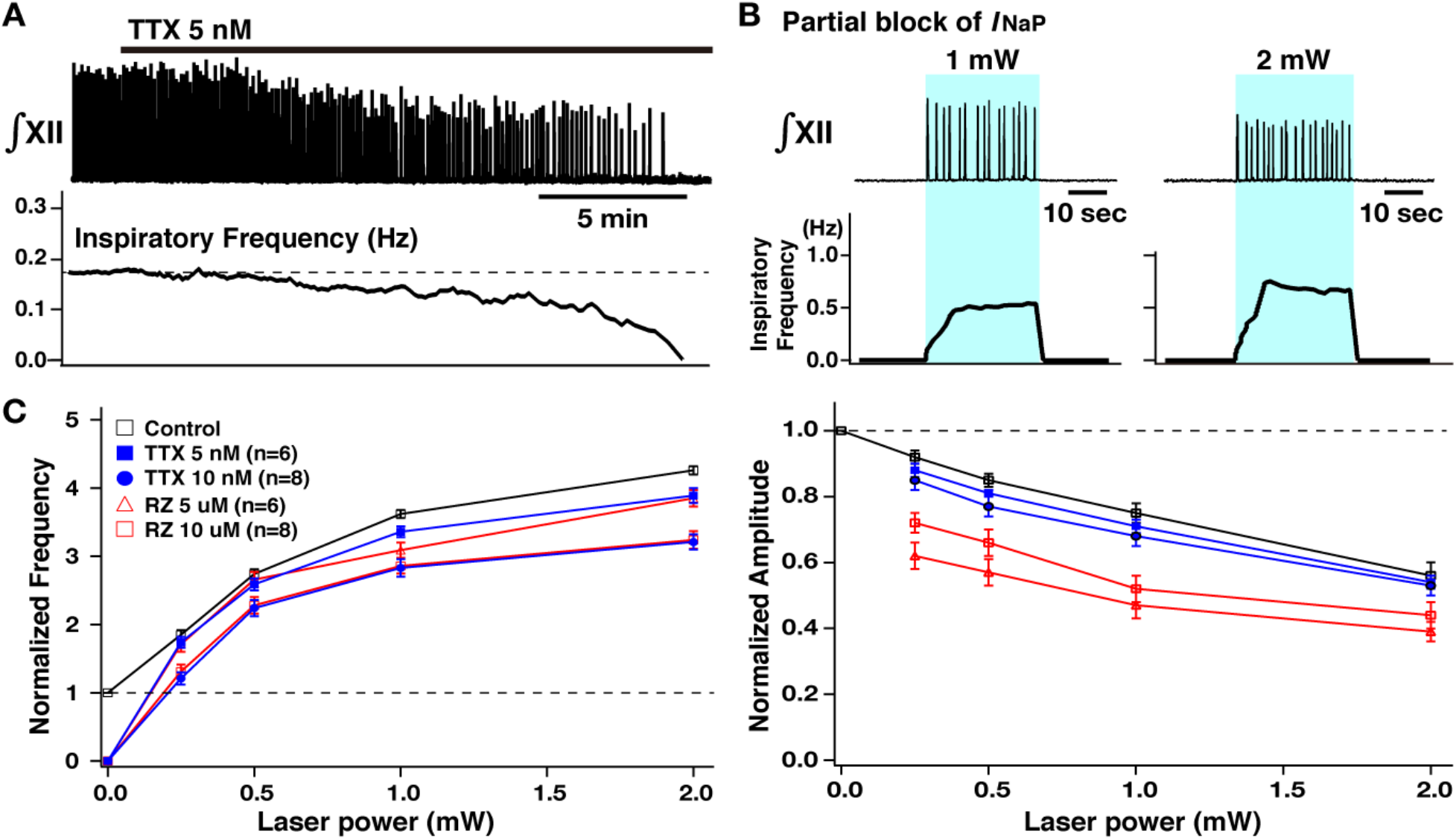
Perturbations of inspiratory burst frequency and amplitude by bilateral preBötC photostimulation during pharmacological block of I_NaP_. (**A**) Example recordings of integrated XII activity with bath application of low concentration of TTX (5 nM), which gradually decreased inspiratory burst frequency, and completely stopped the rhythm within ~25–30 min. The amplitude of XII activity also gradually decreased (~36% before the rhythm stopped in this example). (**B**) Under these conditions of partial block of I_NaP_, bilateral preBötC photostimulation (30 min. after the rhythm stopped, blue shaded epoch) could reinitiate the rhythm, which was also laser power dependent (~202% increase at 1 mW compared to the control, and ~306% increase at 2 mW in the example shown). (**C**) Summary plots (TTX 5nM; n=6, TTX 10 nM; n=8, riluzole (RZ) 5μM; n=6, RZ 10 μM; n=8, mean ± SEM) of the relations between laser power vs normalized inspiratory burst frequency indicates laser power dependent, significant increases of frequency in all cases. The curves for higher concentration of TTX and riluzole are downward-shifted compared to those for the lower concentration as well as those under control conditions (before drug applications). Right panel shows summary plots (TTX 5nM; n=6, TTX 10 nM; n=8, RZ 5μM; n=6, RZ 10 μM; n=8, mean ± SEM) of the relations between laser power vs normalized XII burst amplitude indicating laser power dependent, significant decreases of burst amplitude in all cases. The curves under the I_NaP_ partial block (except for at TTX 5 nM) are downward-shifted compared to those under the control conditions. The decrease in burst amplitudes under RZ is more significant than those under TTX, although there are no significant differences between different concentrations of TTX (blue lines) or RZ (red lines).

Summary data (n = 6 slice preparations at TTX 5nM, n=8 at TTX 10 nM, n=6 at riluzole 5 μM, n=8 at riluzole 10 μM, mean ± SEM) of the relations between normalized inspiratory burst amplitude and laser power presented in Figure 4C (right) show a significant decrease of inspiratory burst amplitude with I_NaP_ attenuation (Wilcoxon signed rank test, p<0.01 at TTX 10 nM and at riluzole 10 μM, and p<0.05 at TTX 5 nM and at riluzole 5 μM). Although significant, changes in amplitude relative to control are very small with TTX block of I_NaP_. The amplitude vs laser power relation with partial I_NaP_ blockade (except for TTX 5 nM) is shifted significantly downward compared to control conditions (TTX 5 nM vs control, p = 0.290; TTX 10 nM vs control, p= 0.036; riluzole 5 μM vs control, p<0.01; riluzole 10 μM vs control, p<0.01 by Wilcoxon signed rank tests), but there were no significant differences (Wilcoxon signed rank test) in the relations for different concentrations of TTX or riluzole (TTX 5 nM vs TTX 10 nM, p=0.229; riluzole 5 μM vs riluzole 10 μM, p = 0.143). The curves obtained under riluzole were more significantly downward-shifted than those under TTX (TTX 5 nM vs riluzole 5 μM, p<0.01; TTX 10 nM vs riluzole 10 μM, p<0.01 by Wilcoxon signed rank tests).

Under conditions of complete block of I_NaP_ (>30 min after either 20 nM TTX or 20 μM riluzole), bilateral preBötC photostimulation could not reinitiate the inspiratory rhythm at any laser power ranging from 0.25 to 5 mW (n=18/18 under TTX, n=11/11 under riluzole), but induced only tonic spiking in preBötC neurons with laser powers >2 mW. This was verified at both the excitatory population activity level (Figure 5), and also the single neuron level. Power spectrum analyses of the recorded tonic activity of the preBötC excitatory population, which was graded with the progressive increases of laser power, verified loss of rhythmic activity at each level of photostimulation-induced tonic activity in comparison to the control activity. In control conditions there is clear definition of spectral peaks for the fundamental and higher harmonic frequencies of the rhythmic population activity, but these features are absent after block of I_NaP_ (Figure 5, n=5). At the cellular level, current-clamp and voltage-clamp recordings confirmed the absence of neuronal rhythmic synaptic drive potentials/currents in preBötC inspiratory glutamatergic neurons (Figure 6).

**Figure 5.**
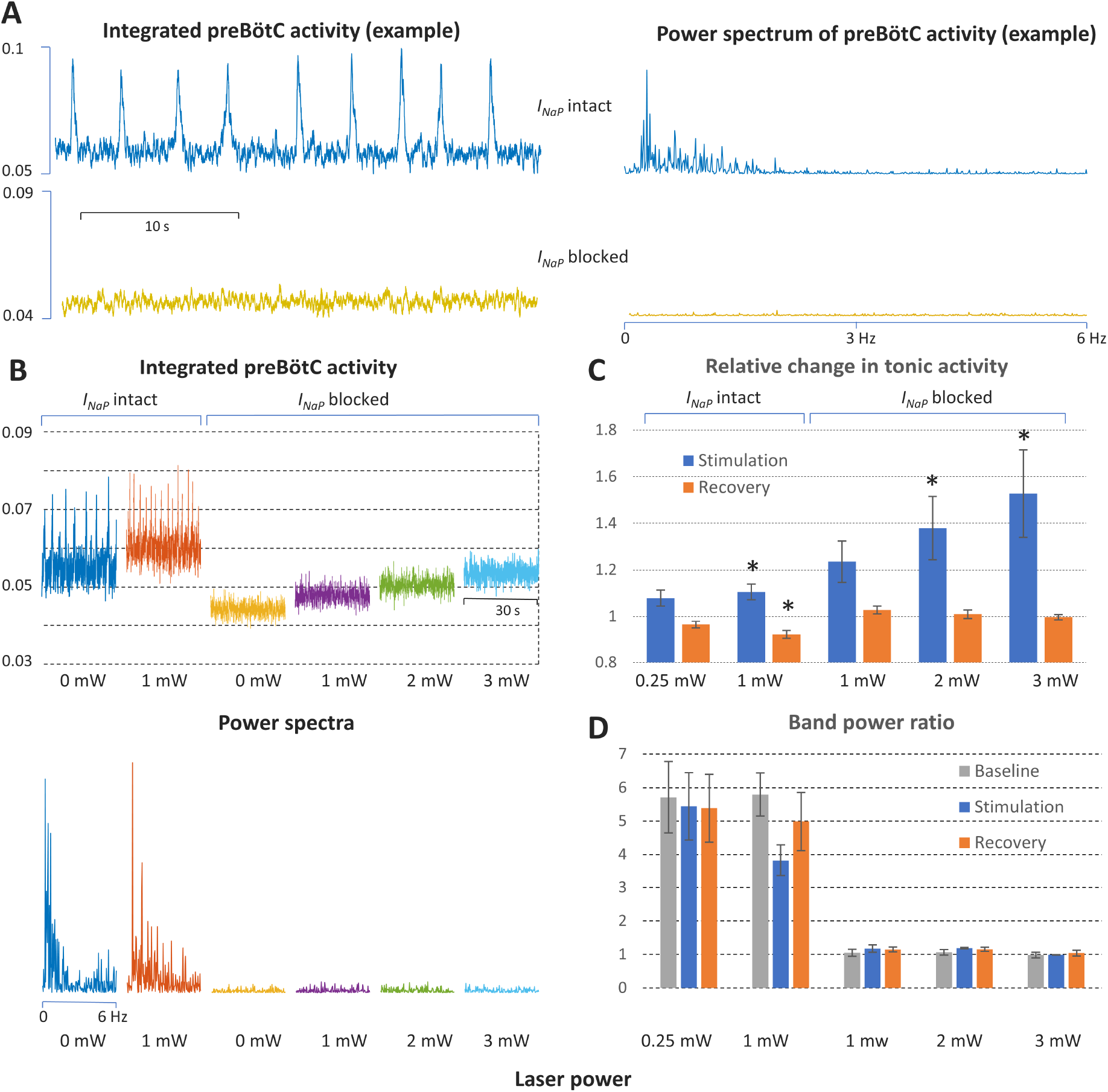
Power spectrum analyses of preBötC neuronal population activity *in vitro* before and after complete block of I_Nap_. (**A**) Examples of preBötC integrated population activity patterns and associated power spectra before and after block of I_Nap_, which eliminates rhythmic preBötC population activity, leaving only a baseline low level of tonic activity or “noise” with a flat power spectrum (lower traces). The power spectra of the rhythmic activity has clear peaks corresponding to the fundamental and higher harmonic frequencies (upper traces). (**B**) Representative experiment illustrating activity patterns and power spectra before and after block of I_NaP_ at various levels of photostimulation (0 and 1 mW in control; 0, 1, 2, and 3 mW after I_NaP_ blocked). With I_NaP_ intact (control conditions, left traces), photostimulation increases the frequency of integrated preBötC inspiratory population activity accompanying an upward shift of the integrated baseline activity due to tonic activity, and there are clear peaks in the power spectra of the rhythmic activity corresponding to the fundamental and higher harmonic frequencies (lower panel). After block of I_NaP_ with 20 nM TTX, there is graded shift in the baseline level of tonic activity during photostimulation, but no rhythmic integrated population activity as indicated by the flat power spectra resembling that of the baseline noise activity in the absence of photostimulation. (**C**) Data from integrated preBötC population recordings showing the change in level of tonic activity relative to baseline during graded photostimulation and the recovery period following stimulation under control conditions with I_NaP_ intact and with I_NaP_ blocked (20 nM TTX). Increasing the level of photostimulation (0.25, 1mW in control; 1, 2, 3 mW with I_NaP_ blocked) in both conditions increases the level of tonic activity. The activity returns to the baseline level (I_NaP_ blocked) or is slightly depressed (control conditions) in the post-photostimulation recovery period. Bars indicate mean ± SEM from 5 slices. * indicates statistically significant difference from unity (p<0.05 by 2-tailed t-test). (**D**) Quantification of the oscillatory component in the preBötC activity before and after complete block of I_NaP_ at different levels of photostimulation as the ratio of the band power in the (0 – 3) Hz frequency range over the band power in the (3 – 6) Hz range. The ratios are high in control conditions, reflecting significantly higher spectral power content in the low frequency band (oscillations) compared to one in the high frequency band (noise) when there is rhythmic activity at baseline, with photostimulation, and during recovery. The ratios are not different and have a value of unity when I_NaP_ is blocked, reflecting the flat power spectra under this condition. Bars indicate mean ± SEM from 5 slices.

**Figure 6.**
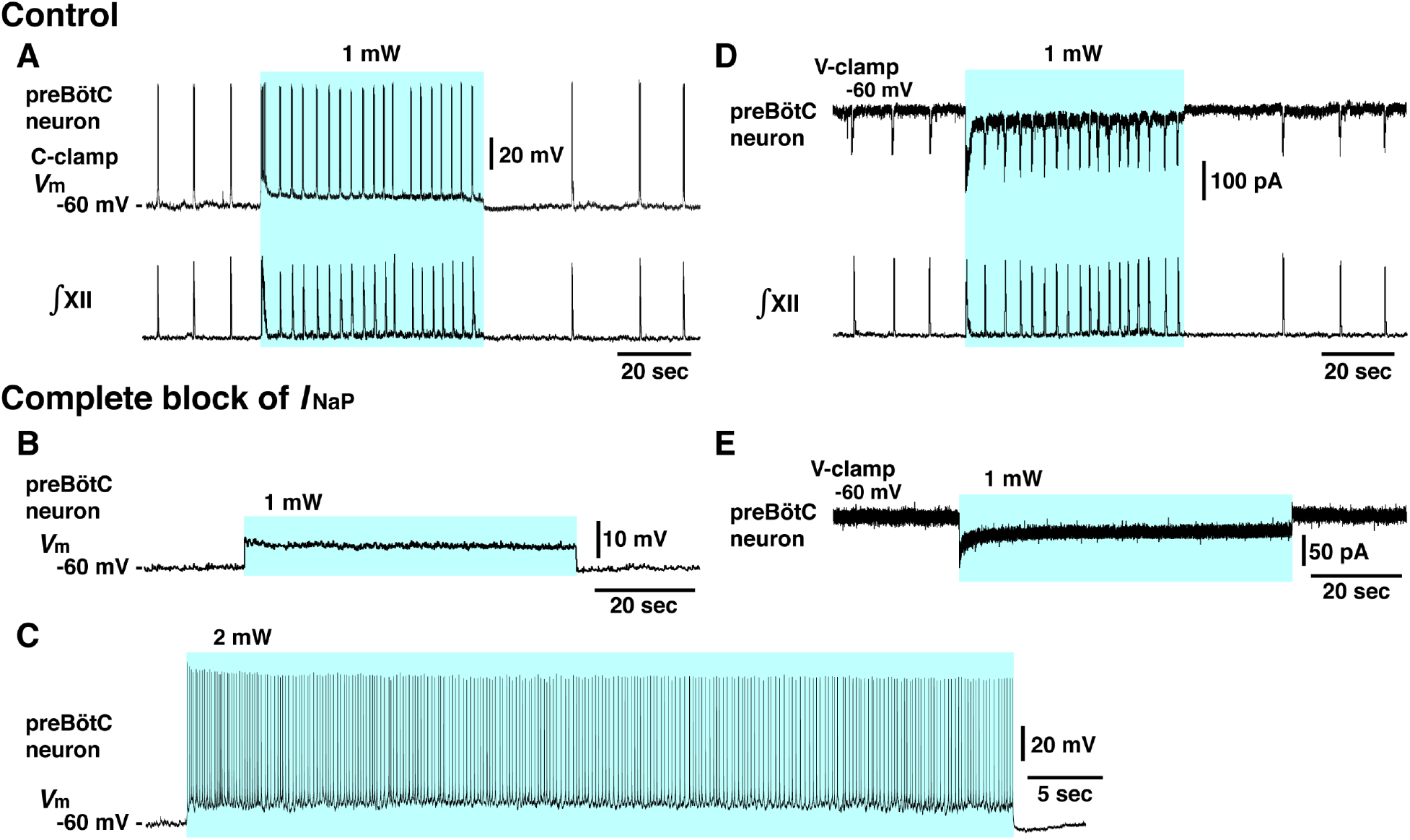
Elimination of inspiratory rhythm at the cellular level after complete block of I_NaP_ *in vitro*. (**A**) Whole-cell current-clamp recordings from td-tomato labeled preBötC inspiratory neuron illustrating rhythmic bursting synchronized with inspiratory hypoglossal motor activity. In control conditions, optogenetic stimulation (1 mW) induced neuronal membrane depolarization (~6 mV) along with a significant increase of inspiratory bursting frequency synchronized with the network bursting frequency indicated by the integrated inspiratory XII activity (∫XII). (**B, C**) With complete block of I_NaP_ (20 nM TTX), photostimulation did not induce rhythmic activity in this neuron, only membrane depolarization without rhythmic synaptic drive potentials or bursting at 1mW laser application (**B**) and only tonic neuronal spiking at higher laser power of 2 mW (**C**). The latter indicates that the neuron retained spiking capabilities at the low concentration of TTX employed, which did not interfere with action potential generation by transient Na+ channels while I_NaP_ was completely blocked (see Supplemental Figure 1). (**D**) Voltage-clamp recordings from td-tomato labeled preBötC inspiratory neuron showing inward rhythmic synaptic drive currents synchronized with inspiratory hypoglossal activities. Under control conditions, optogenetic stimulation (1 mW) induced inward currents and rhythmic synaptic drive currents synchronized with the higher frequency inspiratory hypoglossal activities. (**E**) Rhythmic inspiratory synaptic drive currents did not occur with photostimulation (1 mW) after complete block of I_NaP_ at 20 nM TTX, indicating loss of synaptic interactions and network rhythmic activity.

These results demonstrate that I_NaP_ in preBötC glutamatergic neurons is critically involved in generating the inspiratory rhythm with a voltage-dependent frequency control mechanism *in vitro*.

#### Regulation of inspiratory burst frequency and amplitude by I_CAN_/TRPM4 in preBötC glutamatergic neurons

We next investigated contributions of I_CAN_/TRPM4 to the voltage-dependent control of XII inspiratory burst frequency and amplitude for comparisons to model predictions. We analyzed perturbations of burst frequency and amplitude by bilateral preBötC photostimulation (as described above) after pharmacologically blocking TRPM4 with a specific inhibitor of TRPM4 channels (9-phenanthrol, a putative blocker of I_CAN_). Bath-application of 9-phenanthrol at a concentration proposed to be selective for TRPM4 (50 μM) to the rhythmically active slice preparations from VgluT2-tdTomato-ChR2 neonatal mice (Figure 7A) significantly reduced the amplitude of inspiratory XII bursts (43 ± 4% reduction, n=8, p<0.01 by one sample *t*-test), with little effects on burst frequency (1 ± 1% increase, n=8, one sample *t*-test p = 0.423) in these slices, as we have previously described (Koizumi *et al*., 2018). Under these TRPM4 block conditions (steady-state at >60 min after 9-phenanthrol application, Figure 7C), photostimulation caused a significant increase of inspiratory burst frequency compared to control conditions at a given laser power (0.25 – 2.0 mW, Figure 7D black lines, one sample *t*-test, p<0.01 in all cases), accompanied by a significant laser-power dependent decrease of burst amplitude compared to control (Figure 7D red lines, one sample *t*-test, p<0.01 in all cases). The relations between normalized inspiratory burst frequency and amplitude vs laser power summarized in Figure 7D (n=8 slice preparations, mean ± SEM) show that TRPM4 block significantly shifts the frequency tuning curve upward compared to control conditions (black lines in Figure 7D, Wilcoxon signed rank test, p<0.01). The relation between burst amplitude and laser power is shifted downward significantly compared to control conditions (red lines in Figure 7D, Wilcoxon signed rank test, p<0.01). Note that under baseline conditions of excitability, TRPM4 block does not alter burst frequency, and the upward shift in the frequency tuning curve is opposite to the shift seen with TTX or RZ block of I_NaP_. These findings are consistent with model predictions presented in Figure 1.

**Figure 7.**
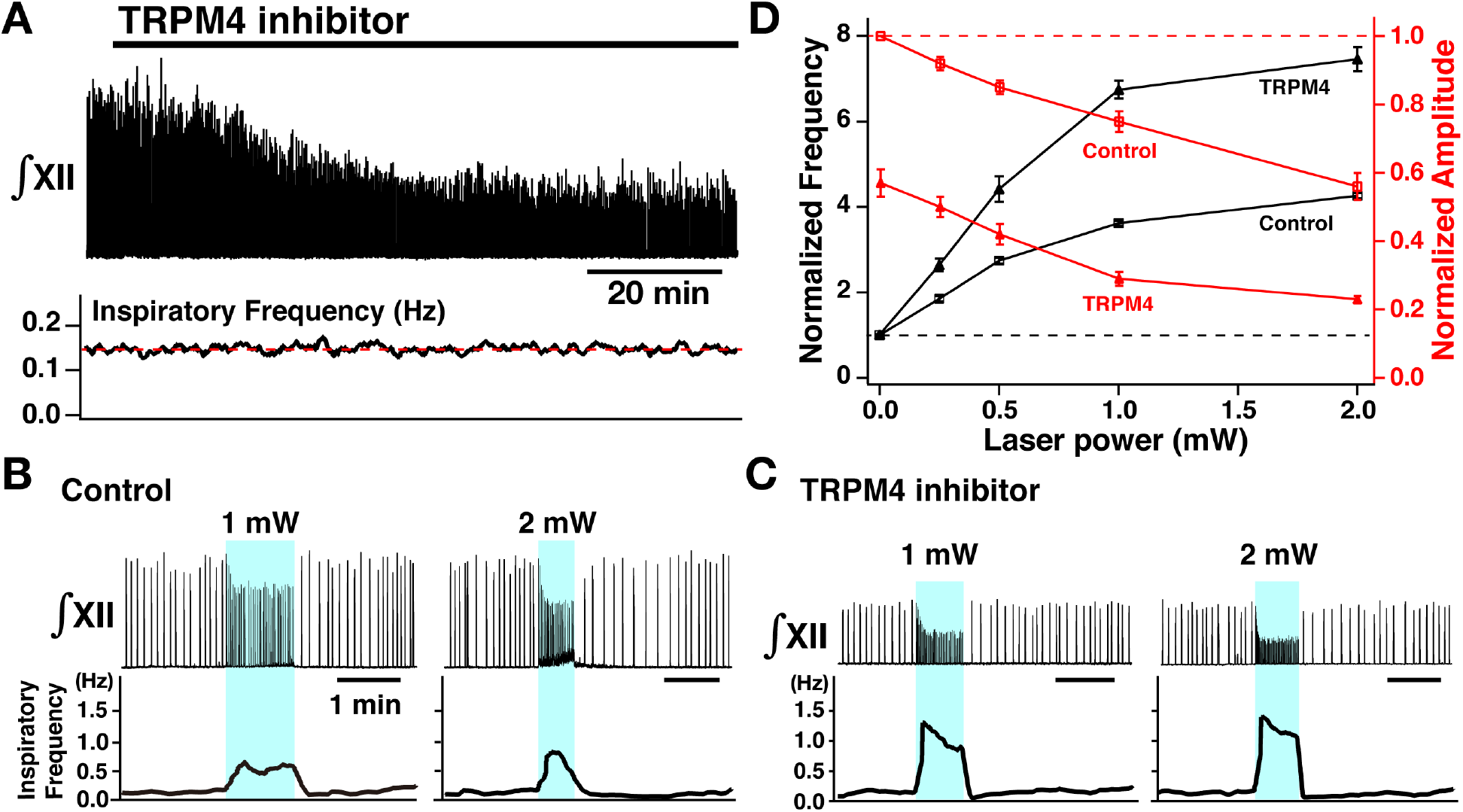
Perturbations of inspiratory frequency and burst amplitude by bilateral photostimulation of the preBötC during TRPM4 pharmacological blockade. (**A**) Upper trace illustrates the time course of integrated XII inspiratory activity during bath application of the specific pharmacological inhibitor of TRPM4 channels (9-phenanthrol, 50 μM), which gradually decreased inspiratory burst amplitude (~58% reduction at steady state ~60 min after drug application), but had little effect on the inspiratory frequency (bottom trace). (**B**) Bilateral photostimulation of the preBötC (Control, before pharmacological block) increased inspiratory burst frequency in a laser power dependent manner (~284% increase at 1 mW, ~354% increase at 2mW) and monotonically decreased inspiratory XII burst amplitude (~25% decrease at 1 mW, ~44% decrease at 2mW). (**C**) Under TRPM4 block conditions (>60 min after 9-phenanthrol application), photostimulation significantly increased inspiratory frequency (~586% increase at 1 mW, ~724% increase at 2mW), and decreased XII inspiratory burst amplitude (~72% decrease at 1 mW, ~78% decrease at 2mW). (**D**) Summary plots of monotonic relations between laser power vs normalized inspiratory frequency (black lines, n=8) and normalized amplitude (red, n=8).

#### Model simulations-data comparisons

For comparisons of model predictions and results from the experimental photostimulation and pharmacological manipulations (Figure 8), we incorporated in the original model (Phillips *et al*., 2019) a Channelrhodopsin-2 current (I_ChR2_) to represent I_App_ for the excitatory neuron population and mimic effects of photo-induced neuronal depolarization. Channelrhodopsin-2 was modeled by a four state Markov channel (see Figure 9), based on biophysical representations of this channel in the literature (Williams *et al*., 2013), which we found could be parameterized as described in Material and Methods to closely match our experimental data on relations between laser power and membrane depolarization of preBötC inspiratory glutamatergic neurons (Figure 8A). In agreement with the experimental results (Figure 4), the model simulations show that reducing g_NaP_ from control values decreases the population-level bursting frequency and amplitude at a given level of excitability (laser power). This reduction also alters the voltage-dependent rhythmogenic behavior with a downward shift in the frequency tuning curve such that the frequency dynamic range is reduced, and rhythm generation can be terminated at a sufficiently low level of g_NaP_ under baseline conditions of network excitability. Due to the different proposed mechanisms of pharmacological action of TTX and riluzole on I_NaP_, there is a larger reduction of amplitude with I_NaP_ block by riluzole than by TTX (as also shown in Phillips and Rubin, PLoS CB, 2019), as seen in the experimental data.

**Figure 8.**
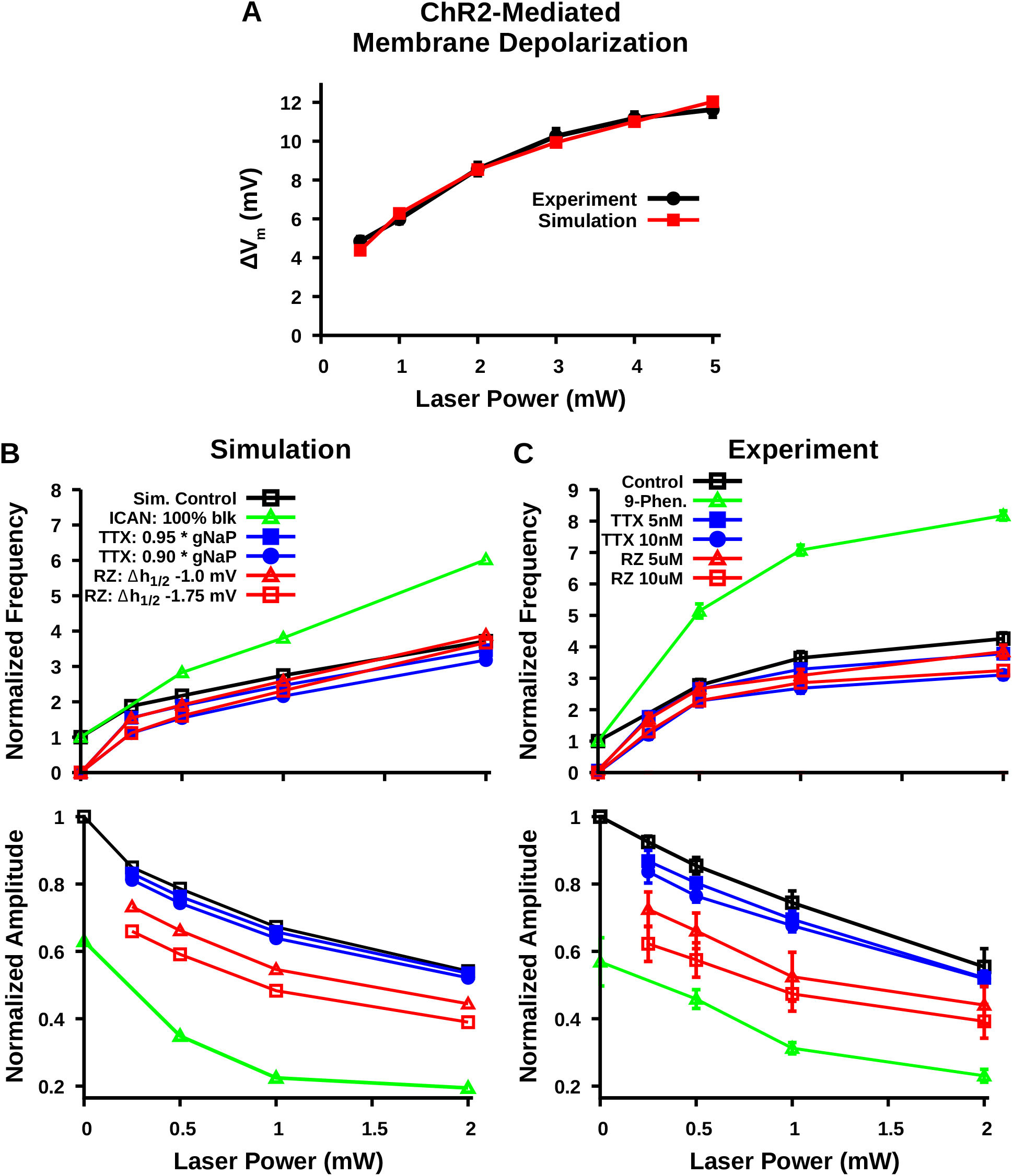
Comparison of experimental and simulated optogenetic photostimulation of the preBötC excitatory network under control conditions and after partial block of I_NaP_ or TRPM4/I_CAN_. **(A)** Matched relationship between neuronal membrane depolarization from baseline (ΔV_M_) as a function of laser power with photostimulation from experimental data (same as shown in Figure 2D) and model simulations. **(B)** Relationship between laser power and burst frequency (top) and amplitude (bottom) of the preBötC network with model simulated blockade of I_NaP_ by TTX and riluzole (RZ) or block of I_CAN_. I_NaP_ block by TTX and I_CAN_ block 9-phenanthrol were simulated by reducing the channel conductances g_NaP_ and g_cAN_, respectively. In contrast, blockade of I_NaP_ by RZ was simulated by a hyperpolarizing shift in the inactivation parameter *V_h_1/2__* and by a partial reduction in *W_max_*; −20% and −25% for simulation of 5μM and 10μM RZ application, as described and justified previously (Phillips and Rubin, 2019). **(C)** Relationship between laser power and inspiratory burst frequency (top) and amplitude (bottom) of the integrated XII output with bath application of the pharmacological blockers of I_NaP_ (TTX or RZ) or I_CAN_ (9-phenanthrol) (same as shown in Figures 4C and 7D). Error bars represent SEM.

Reducing g_CAN_ more strongly decreases population activity amplitude in the model simulations and, in contrast to reducing g_NaP_, has little effect on population bursting frequency under baseline conditions, but strongly augments bursting frequency at higher levels of excitability. This results in an upward shift in the frequency tuning curve that extends the frequency range at the higher levels of population depolarization, consistent with the experimental results (Figure 7). Thus, the model predictions are qualitatively consistent with the experimental data for major features of excitatory network behavior.

## Materials and methods

### Model description and methods

As in the previous paper (Phillips et al., 2019), the preBötC model network is constructed with N=100 synaptically coupled excitatory neurons. Neurons are simulated with a single compartment described in the Hodgkin-Huxley formalism. The addition of Channelrhodopsin-2 current (*I_ChR2_*) is a new feature of the current model compared to Phillips *et al*., 2019. In the updated model, the membrane potential *V_m_* for each neuron is given by the following current balance equation:

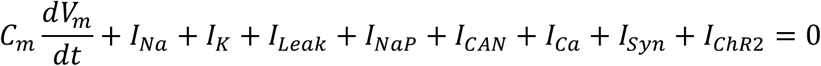

where *C_m_* is the membrane capacitance, *I_Na_, I_K_, I_Leak_, I_NaP_, I_CAN_, I_Ca_, I_Syn_* and *I_ChR2_* are ionic currents through sodium, potassium, leak, persistent sodium, calcium activated non-selective cation, voltage-gated calcium, synaptic and Channelrhodopsin-2 channels, respectively. For a full description of all currents other than *I_ChR2_*, see Phillips *et al*., 2019. The description of *I_ChR2_* was adapted from Williams et al. (2013) and given by the following equation:

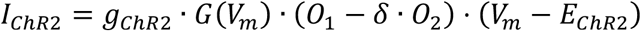

where *g_ChR2_* is the maximal conductance, *O*_1_ and *O*_2_ are light-sensitive gating variables, *δ* is the ratio of the two open-state conductances, *V_m_* is the membrane potential, and *E_ChR2_* is the reversal potential for Channelrhodopsin-2. All parameters given in Table 1. *G*(*V_m_*) is an empirically derived voltage-dependent function given by:

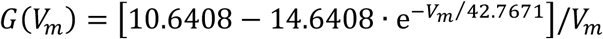

**Table 1.**
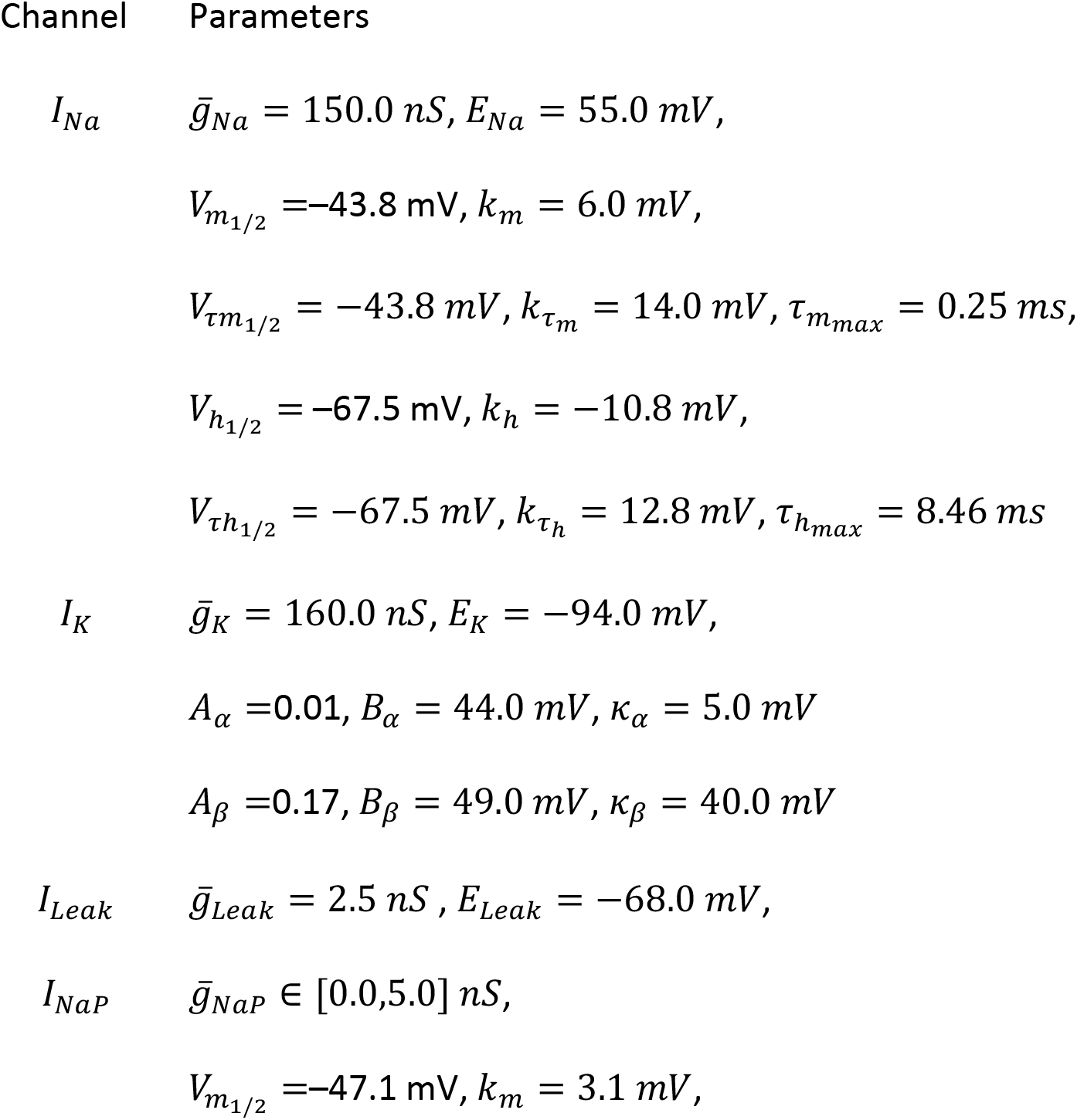
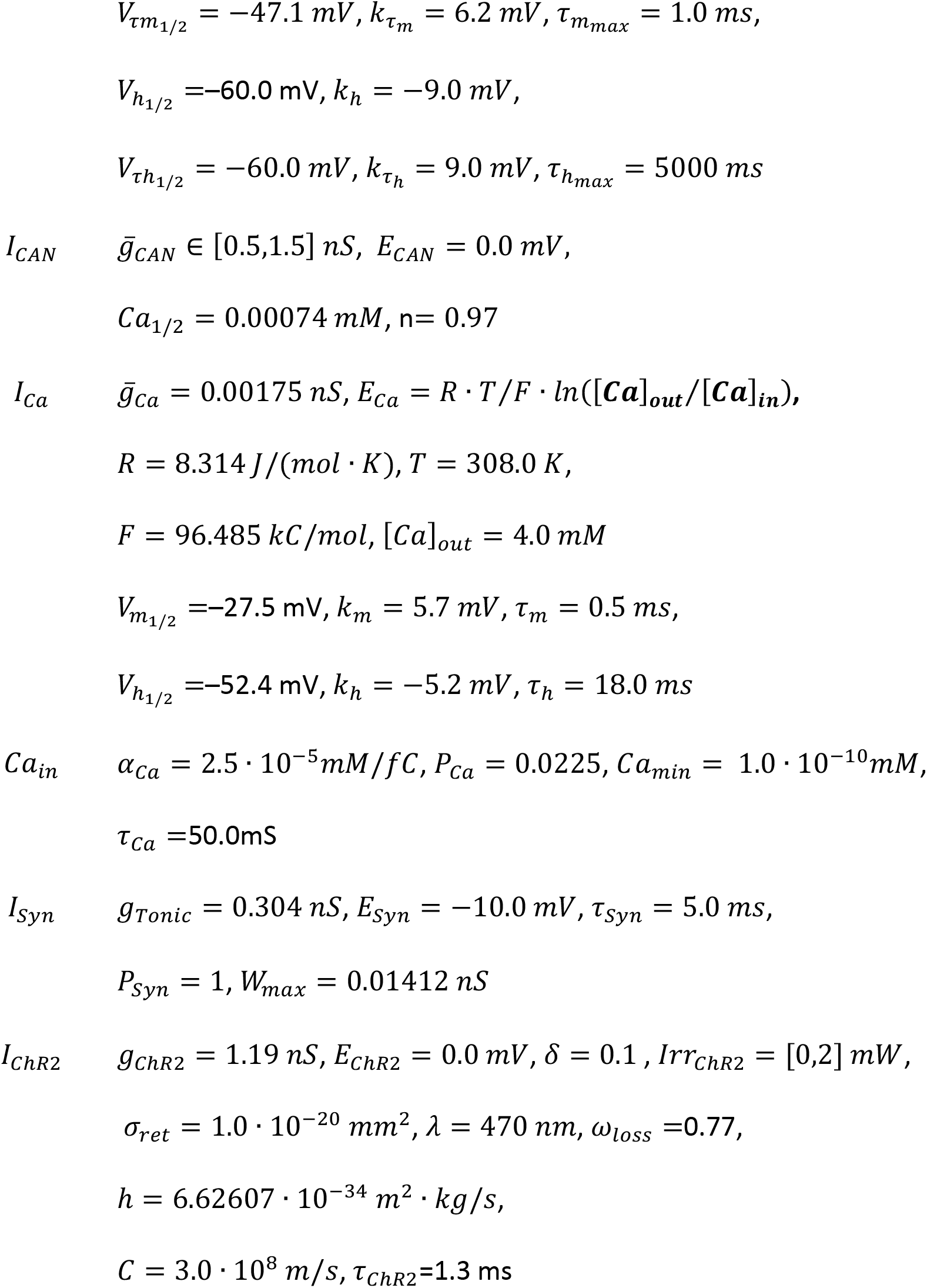
Updated model parameters.

Channelrhodopsin-2 is simulated as a four state Markov channel with two open *O*_1_ and *O*_2_ and two closed states *C*_1_ and *C*_2_ (Figure 9). Transitions from *C*_1_ to *O*_1_ and from *C*_2_ to *O*_2_ are light-sensitive and controlled by the parameter *Irr* which represents optical power with units of milliwatts (mW).

**Figure 9.**
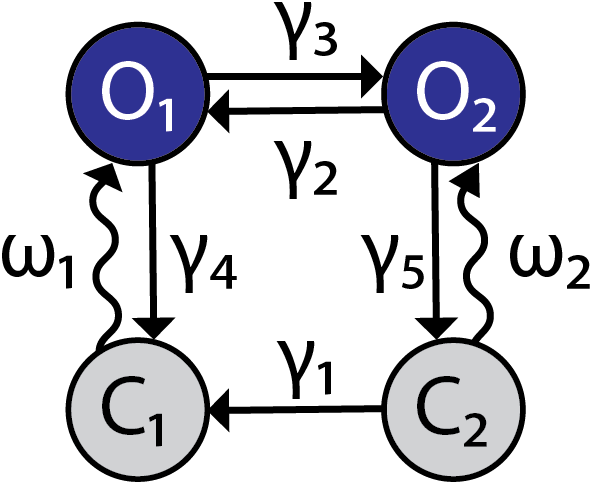
Channelrhodopsin-2 channel configuration. The ChR2 channel activation dynamics are described with a four state Markov model, adapted from (Williams et al., 2013). Transition rates between states are represented by variables *γ*_1–5_ and the light dependent variables *ω*_1,2_.

The rates of change between states are given by the following differential equations:

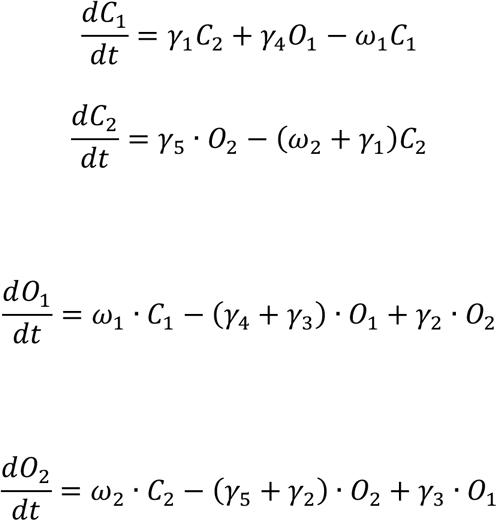

which are consistent with the condition that

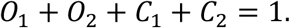

The empirically estimated transition rate (ms^-1^) parameters *ω*_1_, *ω*_2_ *γ*_1_, *γ*_2_, *γ*_1_, *γ*_1_ and *γ*_1_ are defined as follows:

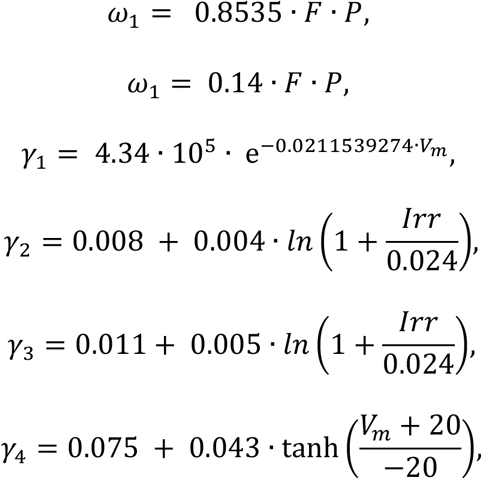

and

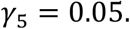

*F* and *P* represent the photon flux (number of photons per molecule per second) and time- and irradiance-dependent activation function for ChR2, respectively. These quantities take the form

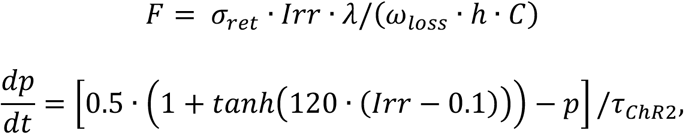

where *σ_ret_* is the absorption cross-section, *λ* is the wavelength of max absorption, *ω_loss_* a scaling factor for loss of photons due to scattering/absorption, *h* is Planks constant, *C* is the speed of light, and *τ_ChR2_* is the time constant of ChR2 activation. Model parameter values are given in Table 1.

### Model Tuning

In order to match experimental data, parameters were slightly modified from the original model tuning in Phillips *et al*. (2019). The updated list of parameters is given in Table 1, where parameter adjustments are indicated with red font.

### Integration methods

All simulations were performed locally on an 8-core Linux-based operating system. Simulation software was custom written in C++. Numerical integration was performed using the exponential Euler method with a fixed step-size (dt) of 0.025 ms. In all simulations, the first 50 s of simulation time was discarded to allow for the decay of any initial condition-dependent transients.

## Experimental Methods

### Animal procedures

All animal procedures were approved by the Animal Care and Use Committee of the National Institute of Neurological Disorders and Stroke.

### Cre-dependent triple transgenic mouse model for optogenetic manipulation of VgluT2-expressing glutamatergic neurons

We have previously established and histologically validated the Cre-dependent double transgenic mouse line (VgluT2-tdTomato) to study population-specific roles of VgluT2-expressing and red fluorescent protein labeled glutamatergic preBötC neurons (Koizumi *et al*., 2016). This VgluT2-tdTomato mouse line was produced using a Slc17a6^tm2(cre)Lowl^/J knock-in Cre-driver mice strain (VgluT2-ires-Cre; IMSR Cat No. JAX: 016963, the Jackson Laboratory, Bar Harbor, ME), in which Cre-recombinase activity is under control by the vesicular glutamate transporter type-2 (VgluT2) promoter (Vong et al., 2011). This line was crossed with a Cre-dependent tdTomato reporter mouse strain [B6;Cg-Gt(ROSA)26Sor^tm9(CAG-tdTomato)Hze^, Rosa-CAG-LSL-tdTomato-WPRE, IMSR Cat No. JAX: 007909, the Jackson Laboratory] to obtain offspring expressing the red fluorescent protein variant tdTomato in Cre-expressing VgluT2-positive (glutamatergic) neurons. The Cre-dependent triple transgenic mouse line VgluT2-tdTomato-ChR2-EFYP, which expresses Channelrhodopsin-2 (ChR2)-EYFP fusion protein and tdTomato reporter fluorescent protein in VgluT2-positive neurons was used for optogenetic photostimulation experiments and in some experiments for imaging glutamatergic neurons for targeted whole-cell patch-clamp electrophysiological recording from these neurons. This transgenic line was obtained by crossing the validated VgluT2-tdTomato line with the Cre-dependent optogenetic mouse strain [B6;129S-Gt(ROSA)26Sor^tm32(CAG-CoP4*H134R/EYFP)Hze^/J, the Jackson Laboratory].

### Rhythmically active medullary slice preparations *in vitro*

We performed optogenetic experiments combined with whole-cell patch-clamp or neural population electrophysiological recordings and pharmacological manipulations in rhythmically active *in vitro* medullary slice preparations (300 – 400 μm thick) from neonatal [postnatal day 3 (P3) to P8] VgluT2-tdTomato-ChR2 mice of either sex. The slice was superfused (4 ml/min) with artificial cerebrospinal fluid (ACSF) in a recording chamber (0.2 ml) mounted on the stage of an upright laser scanning microscope (TCS SP5 II MP, Leica, Buffalo Grove, II). The ACSF contained the following (in mM): 124 NaCl, 25 NaHCO_3_, 3 KCl, 1.5 CaCl_2_, 1.0 MgSO_4_, 0.5 NaH_2_PO_4_, 30 D-glucose equilibrated with 95% O_2_ and 5% CO_2_ (pH = 7.35 – 7.40 at 27 °C). During experiments, rhythmic respiratory network activity was maintained by elevating the superfusate K^+^ concentration to 8 – 9 mM.

### Pharmacological reagents

Tetrodotoxin citrate (TTX) Cat No. 1078, Tocris) at low concentration (5 – 20 nM), riluzole (5–20 μM) (Cat No. R116, Sigma), and 9-phenanthrol (50 uM)— a specific inhibitor of TRPM4 channels at this concentration and putative blocker of I_CAN_ (Cat No. 648492, EMD Millipore Corp., Billerica, MA), were applied to the bathing solution of *in vitro* slice preparations.

### Electrophysiological recordings of preBötC and XII inspiratory activity

We recorded motor outputs in vitro from XII nerve rootlets with fire-polished glass suction electrodes (50 – 100 μm inner diameter) to monitor inspiratory XII motoneuron population activity. In addition, to monitor preBötC inspiratory population activity directly, we performed macro-patch recordings by applying a fire-polished glass pipette (150 – 300 μm inner diameter) directory on the surface of the preBötC region. In some experiments, extracellular neuronal activity was also recorded from preBötC inspiratory neurons with a fine-tipped glass pipette (3 – 5 MΩ resistance) filled with 0.5 M sodium acetate (Sigma) positioned by a computer-controlled 3-dimensional micromanipulator (MC2000, Märzhäuser).

Electrophysiological signals in all cases were amplified (50,000–100,000X; CyberAmp 380, Molecular Devices Union City, CA), band-pass filtered (0.3–2 kHz), digitized (10 kHz) with an AD converter (PowerLab, AD Instruments, Inc., Colorado Springs, CO or Cambridge Electronics Design, Cambridge, UK), and then rectified and integrated digitally with Chart software (AD Instruments) or Spike2 software (Cambridge Electronics Design).

Whole-cell voltage- and current-clamp data were recorded with a HEKA EPC-9 patch-clamp amplifier (HEKA Electronics Inc., Mahone Bay, Nova Scotia, Canada) controlled by PatchMaster software (HEKA; 2.9 kHz low-pass filter, sampled at 100 kHz). Whole-cell recording electrodes (borosilicate glass pipette, 4 – 6 MΩ), positioned with a 3-dimensional micromanipulator (Scientifica, East Sussex, UK), contained the following (in mM): 130.0 K-gluconate, 5.0 Na-gluconate, 3.0 NaCl, 10.0 HEPES, 4.0 Mg-ATP, 0.3 Na-GTP, and 4.0 sodium phosphocreatine, pH 7.3 adjusted with KOH. In all cases, measured potentials were corrected for the liquid junction potential (−10 mV). Series resistance was compensated on-line by ~80%, and the compensation was periodically readjusted.

Whole-cell voltage-clamp recording from optically identified VgluT2-tdTomato expressing inspiratory neurons was used to obtain subthreshold current-voltage (I-V) relationships by applying slow voltage ramps (30 mV/s; −100 to +10 mV) and we computed TTX- and riluzole-sensitive *INaP* by subtracting I-V curves obtained before and after block of I_NaP_ with TTX or riluzole. Data acquisition and analyses were performed with PatchMaster and Igor Pro (Wavemetrics) software (Koizumi and Smith, 2008).

### Optical imaging of preBötC glutamatergic neurons

Optical two-photon live imaging of tdTomato-ChR2-EYFP co-labeled neurons in the slice preparations was performed to verify florescent protein expression and target VgluT2-positive preBötC neurons for patch-clamping. Images were obtained with a Leica multi-photon laser scanning upright microscope (TCS SP5 II MP with DM6000 CFS system, LAS AF software, 20X water-immersion objective, N.A. 1.0, Leica; and 560 nm beam splitter, emission filter 525/50, Semrock). A two-photon Ti:Sapphire pulsed laser (MaiTai, Spectra Physics, Mountain View, CA) was used at 800 – 880 nm with DeepSee predispersion compensation. The laser for two-photon fluorescence imaging was simultaneously used for transmission bright-field illumination to obtain a Dodt gradient contrast structural imaging to provide fluorescence and structural images matched to pixels. These images allowed us to accurately place a patch pipette on the tdTomato-labeled VgluT2-positive neurons to functionally identify preBötC inspiratory glutamatergic neurons, which were active in phase with XII inspiratory network activity.

### Photostimulation of preBötC glutamatergic neuron population

Laser illumination for optogenetic experiments was performed with a blue laser (473 nm; OptoDuet Laser, IkeCool, Los Angeles, CA), and laser power (0.25 - 5.0 mW) was measured with an optical power and energy meter (PM100D, ThorLabs, Newton, NJ). Illumination epochs (sustained 20Hz, 20 msec pulses, variable duration pulse trains) were controlled by a pulse stimulator (Master-8, A.M.P.I., Jerusalem, Israel). Optical fibers, from a bifurcated fiberoptic patch cable, each terminated by an optical cannula (100 μm diameter, ThorLabs), were positioned bilaterally on the surface of the preBötC in the *in vitro* rhythmically active slice preparations.

### Signal analysis of respiratory parameters and statistics

All digitized electrophysiological signals were analyzed off-line to extract respiratory parameters (inspiratory burst frequency and amplitude) from the smoothed integrated XII nerve or preBötC neuronal population inspiratory activities (200 ms window moving average) performed with Chart software (AD Instruments), Spike2 software (Cambridge Electronics Design), and Igor Pro (WaveMetrics) software. Following automated peak detection, burst period was computed to obtain the inspiratory frequency. Inspiratory burst amplitude was calculated as the difference between the signal peak value and the local baseline value. To obtain steady-state perturbations, we analyzed the respiratory parameters during laser illumination excluding the first and last 5-10 sec of laser illumination. Power spectrum analyses were performed using Matlab software. Power spectra were calculated as a squared magnitude of the Fast Fourier Transform of the rectified preBötC activity for 30 sec segments of the recordings during baseline activity, photostimulation and recovery period. The oscillatory component in preBötC activity was quantified as a ratio of the band power of the signal in the 0 – 3 Hz frequency range to the band power of the signal in the 3 – 6 Hz frequency range. For statistical analyses, respiratory parameters during laser illumination were normalized to the control value calculated as an average from ~30 inspiratory bursts before laser illumination. The normality of data was assessed both visually (Quantile vs Quantile Plots) and through the Shapiro-Wilk normality test. Statistical significance (p < 0.01 or p < 0.05) was determined with Student’s *t*-test or Wilcoxon signed rank test (Prism, GraphPad software LLC), and summary data are presented as the mean ± SEM.

## Discussion

In these new combined experimental and modeling studies, we have advanced our understanding of neuronal and circuit biophysical mechanisms generating the rhythm and amplitude of inspiratory activity in the brainstem preBötC inspiratory oscillator in vitro. These studies were designed to further experimentally test predictions of our recent computational model of preBötC excitatory circuits that incorporate rhythm- and amplitude-generating biophysical mechanisms, relying, respectively, on a neuronal subthreshold-activating, slowly inactivating persistent sodium current (I_NaP_), and a calcium-activated non-selective cation current (I_CAN_), mediated by transient receptor potential M4 (TRPM4) channels coupled to intracellular calcium dynamics. Our model explains how the functions of generating the rhythm and amplitude of inspiratory oscillations in preBötC excitatory circuits involve distinct biophysical mechanisms. In essence, the model advances the concepts that (1) a subset of excitatory circuit neurons whose rhythmic bursting is critically dependent on I_NaP_ forms an excitatory neuronal kernel for inspiratory rhythm generation, and (2) excitatory synaptic drive from the rhythmogenic kernel population is critically amplified by I_CAN_ activation in recruited and interconnected preBötC excitatory neurons in the network to generate the amplitude of population activity.

The model predicts that I_NaP_ is critical for rhythm generation and confers voltage-dependent rhythmic burst frequency control, which provides a mechanism for frequency tuning over a wide dynamic range defined by the frequency tuning curve for the rhythmogenic population (i.e., the relationship between applied current/baseline membrane potential across the network and network bursting frequency). Accordingly, reducing neuronal persistent sodium conductance (g_NaP_) decreases the population-level bursting frequency with a weak reduction in amplitude, reduces the frequency dynamic range, and, at sufficiently low levels of g_NaP_, terminates rhythm generation under baseline and other conditions of network excitability *in vitro*, as also shown in Phillips and Rubin, 2019. In contrast, I_CAN_ in the model is essential for generating the amplitude of rhythmic output but not for rhythm generation. As a result, reducing g_CAN_ strongly decreases population activity amplitude and has little effect on population bursting frequency under baseline conditions *in vitro*, but strongly augments bursting frequency, due to a shift in the frequency tuning curve that extends the frequency range, at higher levels of population excitation. These predicted opposing effects of I_NaP_ and I_CAN_ attenuation on the relationship between network excitability and preBötC population rhythmic activity provided a clear basis for model testing, and our experimental results showed an overall strong agreement with the model predictions.

To experimentally test these predictions in the rhythmically active *in vitro* preBötC network in slices from transgenic mice, we used a combination of electrophysiological analyses, pharmacological perturbations of I_NaP_ or I_CAN_, and selective optogenetic manipulations of the preBötC excitatory (glutamatergic) population involved. Our application of optogenetic photostimulation of preBötC glutamatergic neurons to examine the population activity frequency tuning curves under these different conditions of I_NaP_ or I_CAN_ attenuation provided an effective way to probe for underlying mechanisms. We performed new simulations to mimic the optogenetic manipulations under conditions of pharmacological perturbations of I_NaP_ or I_CAN_. The experimental data directionally matched the simulation results, and this correspondence strongly supports the main hypothesis from the model that I_NaP_ and I_CAN_ mechanistically underlie preBötC rhythm and inspiratory burst pattern generation, respectively. The similarities between the model predictions and experimental results support the basic concepts about inspiratory rhythm and burst pattern generation from the model.

### Extensions and limitations of the model

The assumptions and neurobiological simplifications of the original model were previously elaborated in Phillips *et al*. (2019). We have extended the model to allow comparisons of our new experimental data and model simulation results. This included incorporating a Channelrhodopsin-2 current (I_ChR2_) to represent I_App_ for the entire excitatory neuron population and mimic effects of photo-induced neuronal depolarization. We modeled ChR-2 by a four state Markov channel based on channel biophysical representations in the literature (Williams *et al*., 2013). We found that this biophysical model could be parameterized to closely match our experimental data on relations between laser power and membrane depolarization of identified preBötC inspiratory glutamatergic neurons. Furthermore, we found that this channel model parameterized from our cellular-level data yielded frequency tuning curves for the bilateral excitatory population under baseline conditions and with pharmacological perturbations that were in close agreement with the experimentally measured tuning curves. We note that in the model, the assumption is that all excitatory network neurons have the same level of depolarization at a given laser power. In the experimental slice preparations, this may not occur if the neuronal expression levels of ChR2 are nonuniform. We have used transgenic strategies in mice to drive neuronal expression of ChR2 throughout the population of glutamatergic neurons, which has the advantage that there should be efficiency in coverage of this population in terms of channel expression. In support of this assumption, our single neuron electrophysiological data show a small SEM in the relations between neuronal depolarization and laser power, but it is not possible to fully quantify the uniformity of ChR2 expression levels across the entire population of glutamatergic inspiratory neurons.

To compare the model behavior with data from the pharmacological manipulations, we simulated blockade of I_NaP_ by TTX or I_CAN_ inhibition by 9-phenanthrol by reducing the channel conductances g_NaP_ and g_CAN_, respectively. Blockade of I_NaP_ by riluzole (RZ) was simulated by a hyperpolarizing shift in the inactivation parameter *V_h_1/2__* and by a partial reduction in *W_max_* (−20% and −25% for simulation of 5μM and 10μM RZ), as described and justified previously (Phillips and Rubin, 2019) to take into account known or proposed pharmacological actions of RZ from the literature. With this approach, the simulation results were directionally consistent with the data showing that I_NaP_ attenuation, by either low concentrations of TTX or by RZ, and I_CAN_ blockade have opposing effects on the relationship between network excitability controlled by photostimulation and preBötC population burst frequency. We verified the presence of I_NaP_ in preBötC glutamatergic inspiratory neurons from voltage-clamp measurements, and demonstrated partial and complete block of this current by TTX or RZ under our *in vitro* experimental conditions. A difference between the modeling and experimental results that may represent a limitation of the model is that the model network is more sensitive to I_NaP_ block in terms of rhythm perturbation than suggested by the experimental results. In both model simulations and experiments, partial or complete block of this current can terminate rhythm generation; however, rhythm generation can stop at a smaller percentage reduction in I_NaP_ in the model, where it is assumed that I_NaP_ in all network neurons is uniformly reduced by the same amount at a given simulated percent block. This uniformity may not be the case in the *in vitro* network, where heterogeneity of cellular biophysical properties and spatial variability of pharmacological actions associated with drug penetration problems in the slice could cause nonuniformity of block at a given drug concentration.

Furthermore, the biophysical and pharmacological properties of I_NaP_ in preBötC inspiratory neurons are not fully characterized. Our recent biophysical characterization of I_NaP_ for these neurons has confirmed that this current is slowly inactivating (Yamanishi *et al*., 2018), which is a basic assumption of the present and previous models (Phillips *et al*., 2019), although the details of the voltage-dependent kinetic properties are more complex than represented by the first-order kinetics in the present and previous models. Moreover, some additional currents or dynamical processes (e.g., inhibitory currents from local circuit connections, synaptic depression) (Thoby-Brisson *et al*., 2000; Hayes *et al*., 2008; Andrew and Del Negro, 2015; Phillips *et al*., 2018; Baertsch *et al*., 2018; Juárez-Vidales, *et al*., 2021; Abdulla *et al*., 2021; Revill *et al*., 2021), which are not considered in the model, may play a role in augmenting/shaping inspiratory bursting. Nonetheless, with the kinetic and pharmacological properties incorporated, the directional trends in the experimental results for the various pharmacological manipulations of this channel are correctly predicted by the model simulations.

### Further insights into mechanisms of rhythm and amplitude generation in preBötC excitatory circuits in vitro

Many previous experimental and theoretical analyses have focused on the rhythmogenic mechanisms operating under *in vitro* conditions in rhythmically active slices to provide insight into biophysical and circuit processes involved, which as discussed in a number of reviews, has potential relevance for rhythm generation *in vivo* (Feldman and Del Negro, 2006; Richter and Smith, 2014; Phillips *et al*., 2019). There is agreement that preBötC circuits have intrinsic autorhythmic properties, particularly because when isolated in slices *in vitro*, these circuits continue under appropriate conditions of excitability to generate rhythmic activity that drives behaviorally-relevant inspiratory hypoglossal motoneuronal output. However, there is currently no consensus about the underlying rhythmogenic mechanisms. The endogenous rhythmic activity *in vitro* has been suggested to arise from various cellular and circuit biophysical mechanisms including from a subset of intrinsically bursting neurons which, through excitatory synaptic interactions, recruit and synchronize neurons within the network (pacemaker-network models) (Rekling and Feldman, 1998; Toporikova and Butera, 2011; Ramirez, Tryba and Pena, 2004), or as an emergent network property involving recurrent excitation (Jasinski *et al*., 2013) and/or synaptic depression (group pacemaker model) (Rubin *et al*., 2009a). More recent “burstlet” models for rhythm generation emphasize how rhythm emerges in low excitability states due to synchronization of subsets of excitatory bursting neurons (Del Negro, Funk and Feldman, 2018), but do not fully detail the putative cellular-level or other biophysical mechanics underlying rhythm and amplitude generation.

The present experimental results are consistent with previous findings indicating that the subthreshold-activating, slowly-inactivating I_NaP_ is a critical neuronal conductance for inspiratory rhythm generation and neuronal bursting in vitro (Koizumi and Smith, 2008; Yamanishi *et al*., 2018). As our previous experimental and modeling results have indicated, the voltage-dependent and kinetic properties of these channels are fully capable of orchestrating rhythmic bursting at neuronal (e.g., Yamanishi *et al*., 2018) and excitatory neuron population levels (Koizumi *et al*., 2016). In the experiments blocking I_NaP_, we tested for non-I_NaP_ dependent rhythmogenic mechanisms after fully blocking this conductance as verified by our cellular-level measurements. These experiments were designed to determine if there are additional emergent rhythmogenic mechanisms inherent within the excitatory network at various levels of excitation, controlled in our experiments by sustained bilateral photostimulation that could generate graded levels of tonic population activity in the network. We did not find any coherent rhythmic population activity under these conditions that would indicate the presence of other important rhythmogenic mechanisms capable of producing rhythmic population bursting on any time scale, which is consistent with our model. These results do not support the concept that rhythm generation under *in vitro* conditions is an emergent network property through recurrent excitation (Jasinski *et al*., 2013) and/or synaptic depression (group pacemaker model) (Rubin *et al*., 2009a). Such emergent rhythms are theoretically possible as shown by previous modeling studies (Jasinski *et al*., 2013; Rubin *et al*., 2009a; Guerrier *et al*., 2015), but the regenerative population burst-generating and burst-terminating mechanisms incorporated in these models are apparently not sufficiently expressed in the *in vitro* preBötC network.

We note that our new measurements indicate that while I_CAN_/TRPM4 activation is not involved in rhythm generation under baseline conditions of network excitability, since inhibiting this current does not significantly affect the rhythm, we also show that reducing this current does shift the frequency tuning curve at higher levels of network excitation. This occurs because activation of this current augments excitatory synaptic interactions, which can affect network bursting frequency. As discussed above, these observed effects of reducing I_CAN_/TRPM4 on bursting frequency at elevated levels of network excitation, which we were able to reveal with our photostimulation paradigm, is a basic feature and entirely consistent with predictions of our model.

Our new experimental results confirm our previous experimental and modeling results indicating that endogenous activation of I_CAN_/TRPM4 is critically involved in generating the amplitude of population activity. The correspondence between the experimental data and model predictions supports the concept in the model that activation of I_CAN_ is largely due to synaptically-activated sources of neuronal Ca^2+^ flux such that I_CAN_ contributes to the excitatory inspiratory drive potential and regulates inspiratory burst amplitude by augmenting the excitatory synaptic current. Our data also show that I_NaP_ is involved, to a small degree, in generating the amplitude of rhythmic population activity at a given level of excitatory network excitation. This is due to the subthreshold activation of I_NaP_ and its voltage-dependent amplification of synaptic drive; indeed, application of riluzole, which is thought to impact synaptic transmission, affected population amplitude much more than application of TTX (Figure 4, Figure 8). This contribution to amplitude is smaller than that from I_CAN_/TRPM4 activation, however. In general, our results support the concept from our original model that I_CAN_ activation in a subpopulation of preBötC excitatory neurons is critically involved in amplifying synaptic drive from a subset of neurons whose rhythmic bursting is critically dependent on *I_NaP_*. This later subpopulation forms the kernel for rhythm generation *in vitro*.

### Previous pharmacological studies and proposed roles of I_NaP_ in preBötC inspiratory network rhythm generation

The role of I_NaP_ in rhythm generation within preBötC circuits is highly debated in the field with diverse and contradictory experimental results (Del Negro, Morgado-Valle and Feldman, 2002; Del Negro *et al*., 2005; Pena *et al*., 2004; Ramirez, Tryba and Pena, 2004; Feldman and Del Negro, 2006; Smith *et al*., 2007; Pace *et al*., 2007; Koizumi and Smith, 2008; Ashhad and Feldman, 2020). Pharmacological studies (Smith *et al*., 2007; Koizumi and Smith, 2008) using RZ or low concentrations of TTX to block I_NaP_ in preBötC neurons/circuits, as in the present studies, demonstrated a large reduction of preBötC inspiratory bursting frequency at cellular and network levels, with relatively smaller reductions of inspiratory network activity amplitude; fully blocking I_NaP_ completely eliminated inspiratory rhythm generation within isolated preBötC circuits. These experimental results are consistent with the computational hypothesis presented here and earlier that I_NaP_ is an essential biophysical component of inspiratory rhythm generation (Butera, Rinzel and Smith, 1999a; Butera, Rinzel and Smith, 1999b; Smith *et al*., 2000; Phillips *et al*., 2019). Furthermore, another experimental result (Koizumi and Smith, 2008), similar to our present photostimulation results, was that preBötC network excitation (e.g., by the known preBötC neuronal excitant Substance-P) could reinitiate the inspiratory rhythm only with partial block of I_NaP_, but not with complete block of I_NaP_ after the rhythm was terminated by RZ or TTX.

In contrast to the above studies, other experimental studies *in vitro* (Del Negro, Morgado-Valle and Feldman, 2002; Pena *et al*., 2004; Pace *et al*., 2007) showed that pharmacological block of I_NaP_ with bath application or microinjection into the preBötC of RZ or TTX reduced the amplitude of inspiratory network activity significantly, but caused little or no perturbations of inspiratory burst frequency. Accordingly, the conclusion was that I_NaP_ does not play an important role in inspiratory rhythm generation in preBötC circuits *in vitro*. Alternative hypotheses proposed are that I_NaP_-dependent and I_CAN_-dependent bursting mechanisms collectively contribute to inspiratory rhythm generation (Pena *et al*., 2004), or an emergent network rhythm hypothesis—the group pacemaker model— (Rekling and Feldman, 1998; Feldman and Del Negro, 2006)(Rekling and Feldman, 1998; Feldman and Del Negro, 2006), and lately the burstlet hypothesis (Kam *et al*., 2013; Del Negro, Funk and Feldman, 2018; Ashhad and Feldman, 2020) for inspiratory rhythm generation that does not involve I_NaP_ as a fundamental burstlet (rhythm) or burst (pattern) generating mechanism. Interestingly, in a study by Pace *et al*. (2007), preBötC network excitation by Substance-P could revive the inspiratory rhythm after cessation caused by pharmacological block of both I_NaP_ and I_CAN_. They suggested that block of I_NaP_ affects the state of neuronal excitability of the preBötC, but does not cause a fundamental breakdown in rhythmogenic mechanisms *in vitro*.

The contradictory findings in these different experimental studies can be explained and predicted by computational modeling by considering non-uniformity of I_NaP_ blockade, in vitro slice thickness, and the order in which neurons (patterning vs rhythmogenic neurons) in the network are affected (Phillips and Rubin, 2019). Penetration of bath applied I_NaP_ blockers in in vitro slice preparations depends on passive diffusion. Therefore, the magnitude and progression of I_NaP_ block across the slice will not be uniform and may not completely penetrate thicker slices. Moreover, the preBötC network dynamically expands/contracts in the rostralcaudal axis (Baertsch *et al*., 2019), which suggests that rhythmogenic neurons are preferentially located near the center of a larger population of pattern forming inspiratory neurons. Therefore, I_NaP_ block in thick slices is predicted to primarily affect the amplitude of the inspiratory rhythm rather than frequency and rhythm generation. Consistent with model predictions, previous studies which found that pharmacological block of I_NaP_ reduces amplitude without affecting frequency (Del Negro, Morgado-Valle and Feldman, 2002; Pena *et al*., 2004; Pace *et al*., 2007) all use thick *in vitro* slices. In addition, thicker slices than used in our studies contain not only the preBötC, but also other network components that may introduce additional mechanisms, which is why we use thin slice preparations to allow intrinsic properties of the isolated but functional preBötC circuits to be more readily analyzed *in vitro*. Indeed, with more intact brainstem respiratory pattern generation networks producing the fuller complement of inspiratory and expiratory neuronal activity in mature rodents, I_NaP_ blockers do not disrupt rhythm generation (Smith *et al*., 2007; Rubin *et al*., 2009b; Rubin and Smith, 2019), where neuronal dynamics in the preBötC are controlled by more complex synaptic interactions, including inhibitory circuit interactions, with new rhythmogenic regimes (Rubin *et al*., 2009b; Rubin and Smith, 2019). Interestingly, modeling results suggest there are network states and pharmacological conditions under which the inspiratory rhythm may be highly dependent upon I_NaP_ in the more intact respiratory pattern generation network (Phillips and Rubin, 2019; Rubin and Smith, 2019), and these predictions require further experimental testing.

Regardless of this greater complexity, our present study has confirmed our previous results that I_NaP_, is essential for inspiratory rhythm generation in the isolated preBötC excitatory circuits *in vitro*, representing a major rhythmogenic mechanism in this reduced state of the respiratory network. An important prediction of our model (see Phillips *et al*. (2019) Fig. 9) is that the subpopulation of excitatory neurons with I_NaP_-dependent rhythmogenic properties forming the rhythmogenic kernel may be relatively small compared to the neuronal subpopulation(s) with the I_CAN_-dependent mechanism that critically amplifies the rhythmic drive from the kernel to generate and control the amplitude of excitatory population activity. This prediction also requires direct experimental testing, such as by dynamic multicellular Ca^2+^ imaging in glutamatergic neurons in vitro (Koizumi *et al*., 2018).

### Summary

Our exploitation of optogenetic control of glutamatergic neuron network excitability, in combination with specific pharmacological manipulations of neuronal conductances have enabled rigorous testing of predictions of our previous model of preBötC excitatory circuits. The basic predictions of the model for cellular and network behavior under the experimental conditions tested show a strong overall agreement with our experimental results. This agreement advances our understanding of neuronal and circuit biophysical mechanisms generating the rhythm and amplitude of inspiratory activity in the brainstem preBötC inspiratory oscillator *in vitro* and demonstrates the predictive power of our model.

## Acknowledgements

This work was supported in part by the Intramural Research Program of the National Institutes of Health (NIH), National Institute of Neurological Disorders and Stroke (JS, HK), and grants of NSF DMS 1951095 (JR), NSF DMS 1724240 (JR, RS), NIH R01 AT008632 (YM), NIH U01 EB021960 (YM) and Georgia State University B&B Seed Grant (YM).

---

**Supplementary Figure 1.**
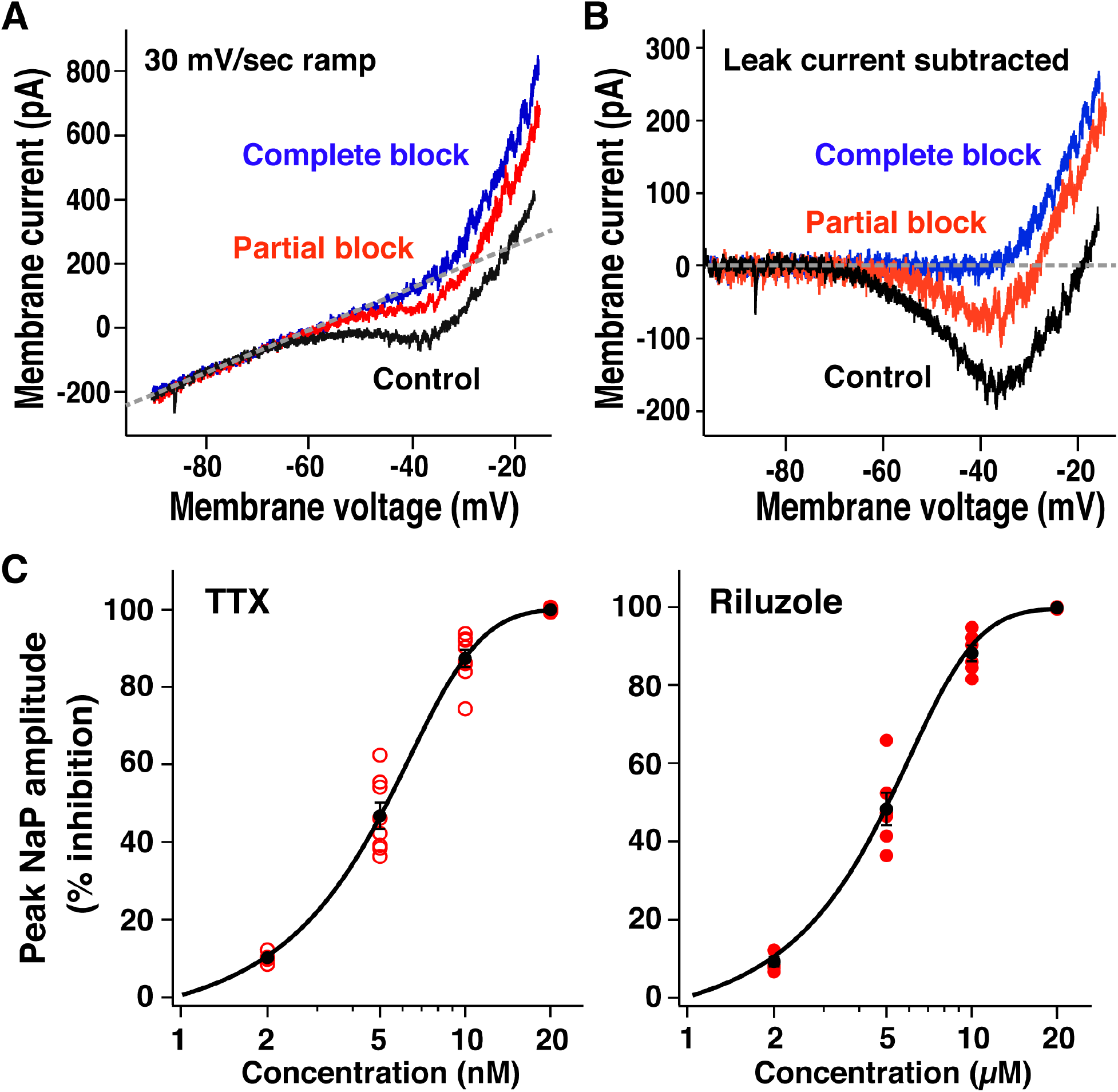
Pharmacological profile of block of I_NaP_ in preBötC inspiratory glutamatergic neurons. (**A**) Example of subthreshold current-voltage (I-V) relationships measured from whole-cell voltage-clamp recording obtained by applying slow voltage ramps (30 mV/s; −100 to +10 mV) from optically identified VgluT2-tdTomato expressing preBötC inspiratory neuron. I-V curves were measured in control, partial block of I_NaP_ (5 nM TTX), and complete block of I_NaP_ (20 nM TTX) conditions. (**B**) TTX-sensitive I_NaP_ obtained from raw whole-cell recordings in **A** by subtracting I-V curves measured before and after application of TTX, illustrating reductions of I_NaP_ inward current with partial and complete block. (**C**) Relations between percent reduction of peak I_NaP_ amplitude (measured at −40 – −35 mV after I-V curve subtraction) and TTX or riluzole concentrations for VgluT2-tdTomato expressing inspiratory neurons. Data points are fitted with a sigmoid curve.

